# Universal barcoded connectomics for massively parallel single-neuron projection mapping in diverse vertebrate species

**DOI:** 10.64898/2026.11.19.767886

**Authors:** Huihui Qi, Xinghaoyun Wan, Julian Cheron, Maryam Majeed, Catherine Elorette, Angela Fan, Zetian Yang, Manjari M-G Anant, Dylan Z. Faltine-Gonzalez, Caleb Shi, Craig Washington, Maggie Lowman, James Bourne, Greg D. Horwitz, Peter H. Rudebeck, Xiaoyin Chen, Michael A. Long, Jon T. Sakata, Justus M Kebschull

## Abstract

Neuroscience relies on diverse model species to understand animal behavior and brain function. Yet, tools for rapid, high-resolution brain connectivity mapping remain unavailable in most vertebrate species, slowing progress in many neuroscience models and rendering cross-species circuit comparisons challenging. Here, we overcome this barrier by establishing sequencing-based barcoded connectomics for rapid single-neuron tracing in seven vertebrate species ranging from frogs to birds to non-human primates. We develop a general solution to viral tropism and barcode trafficking, validate our reagents by mapping the projections of thousands of neurons in the songbird HVC, and discover a single set of reagents that functions across highly divergent species. Together, we provide ready-to-use barcoding reagents for diverse species and a simple framework for further adapting them if necessary, enabling high throughput single-neuron tracing across vertebrates.

## Main

The study of diverse animal species that exhibit unique and tractable behaviors is a central approach to understanding brain function^1–7^. Similarly, the comparison of species with divergent or convergent behaviors is a powerful lever to uncover the computational principles underlying behavior^1,8,9^. While broadly applicable behavioral^10,11^, molecular^12–14^, and genetic^15,16^ tools have recently transformed neuroscience investigations across vertebrate species, circuit mapping tools have fallen behind: In the large brains of most vertebrates, rapid high-resolution and brain-wide circuit mapping is currently infeasible, with limited options in mammals and essentially none in non-mammalian vertebrates. Accordingly, established neuroscience models, such as songbirds and non-human primates, and emerging models, such as amphibians, lack rapid high-resolution circuit tracing tools. As a result, modern high-resolution neuroscience investigations in these species into topics as fundamental as learning and memory, acoustic communication and motor learning, and regeneration and plasticity are strongly impeded by the sparsity of connectivity data.

An ideal circuit mapping tool for use in diverse species should have single neuron resolution, seamlessly integrate connectivity with transcriptomic cell type information of the same cells, and yield comprehensive datasets quickly and at low cost from few animals. The same tool should also work robustly in many species to enable comparison of equivalent connectivity data across species and easy adoption by the community. Barcoded connectomics tools, including MAPseq and its *in situ* sequencing based cousin BARseq, address many of these requirements by enabling rapid, large-scale single-neuron projection mapping with sequencing-based readout of virally delivered barcodes and cell-type information^17–20^. While theoretically applicable to any system, these methods were specifically developed for certain regions of the mouse brain^17,18^. Recent applications of the laboratory mouse reagents to Alston’s singing mice^21^ and macaques^22^ provided important biological insights, but show significantly reduced efficiency, underscoring how difficult it remains to extend mouse connectivity mapping approaches across even closely related mammalian species, let alone more distantly related vertebrate clades such as birds and amphibians.

Overcoming these limitations, here we establish a general platform for single-neuron projection mapping across a diverse set of vertebrates, spanning amphibians, birds, and mammals including non-human primates, using a modular solution to viral tropism and barcode trafficking. We independently address the two barriers to viral tropism, viral entry and expression, through targeted viral pseudotyping and modulating the anti-viral host response. We then engineer a carrier protein that efficiently traffics barcodes in highly divergent vertebrates, identify a single set of reagents that work well across species, and benchmark our reagents in songbird premotor region HVC (proper name) against single-neuron ground truth data. Together, these advances enable the systematic, rapid, and routine mapping and comparison of single-neuron projection patterns across vertebrates.

## Results

### Pseudotyped MAPseq viruses enable neuronal infection in mouse, zebra finch, and Xenopus

A major bottleneck in applying any viral circuit tracing tool, including MAPseq barcoded connectomics, to diverse species is viral tropism, i.e., the (in)ability of a particular virus to transduce a target cell type. MAPseq is based on a replication incompetent Sindbis virus (SINV), an alphavirus capable of driving rapid and robust expression of single RNA barcodes in neurons^17,23^. Although SINV efficiently transduces neurons throughout much of the mouse brain, its tropism is limited: it fails to infect neurons in the mouse cerebellar nuclei (CN) and exhibits poor transduction efficiency in the avian brain (**Fig. 1a**). Decreased infection efficiencies proportionally limit the number of cells that can be traced per experiment, often preventing the robust mapping of connectivity.

**Figure 1.**
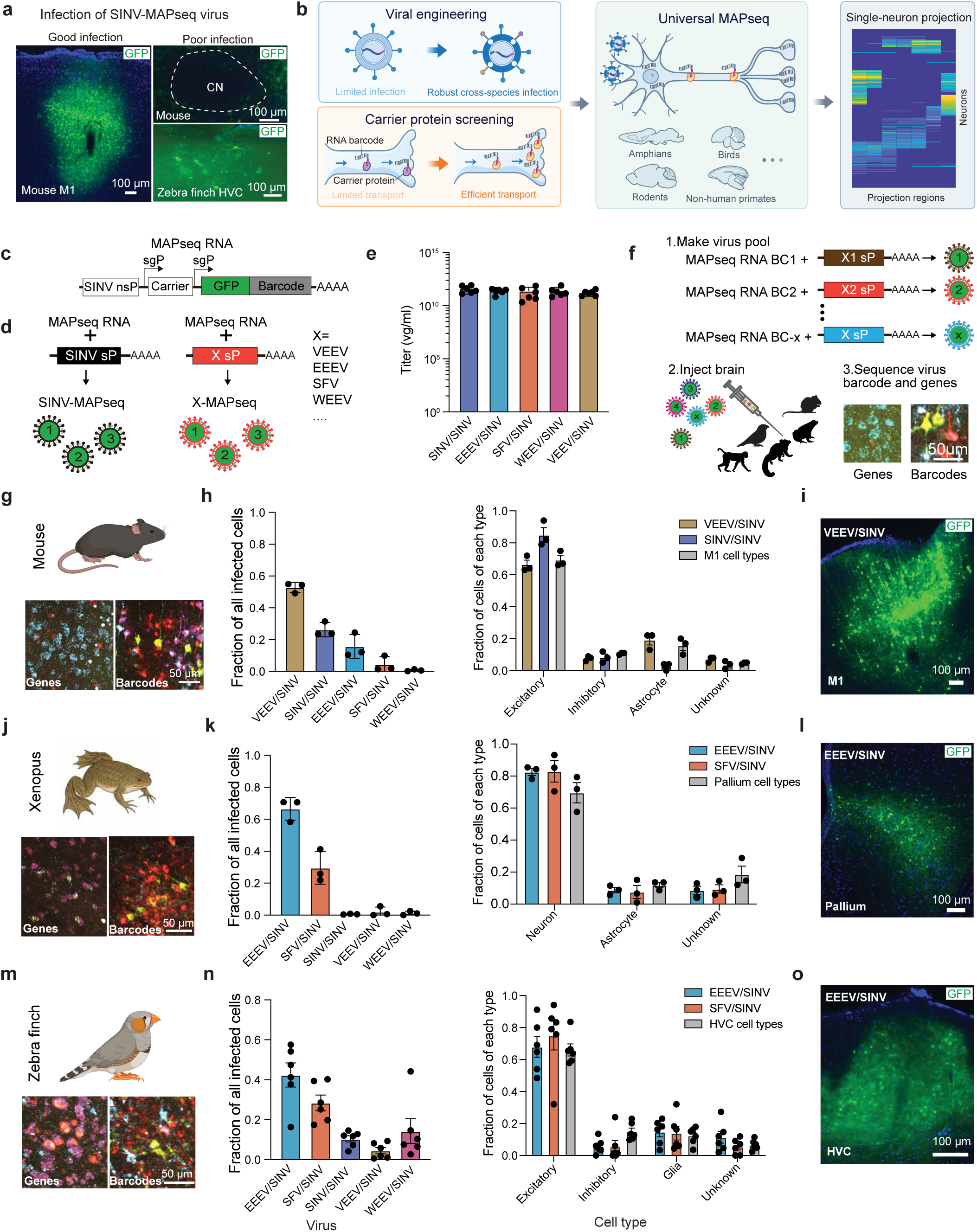
Developing a MAPseq virus for mice, Xenopus, and zebra finches. (a) The original MAPseq virus efficiently infects mouse primary motor cortex (M1) but shows poor infection in the mouse cerebellar nuclei (CN) and zebra finch HVC. (b) Schematic of the universal MAPseq strategy for single-neuron projection mapping in diverse species. (c) Genome structure of the original MAPseq virus. (d) Pseudotyping leaves the viral genome unchanged while varying the viral structural proteins and thus the virus’ infective properties. (e) Titers of MAPseq viruses packaged with the structural proteins of different alphaviruses, showing high titers independent of the structural protein donating virus. (n = 6) (f) Strategy for virus screening. Each pseudotyped virus is assigned a unique identifier barcode, allowing simultaneous identification of viral identity and cellular gene expression by BARseq *in situ* sequencing of infected cells. (g-o) Screening and validation of pseudotyped viruses in mouse M1, *Xenopus tropicalis* pallium, and zebra finch HVC. Representative *in situ* sequencing images showing viral barcodes and gene expression 24 h after infection in mouse (g), Xenopus (j), and zebra finch (m). In mouse M1, VEEV/SINV and SINV/SINV produced robust infection, with VEEV/SINV infecting a slightly higher proportion of astrocytes than SINV/SINV (h). Independent validation confirmed robust infection by VEEV/SINV in mouse M1 (i). EEEV/SINV and SFV/SINV were the two top-performing viruses identified in Xenopus and zebra finch, and showed cell-type distributions similar to those of the injected brain regions, suggesting broad cellular tropism (k, n). Independent validation confirmed robust infection with EEEV/SINV in Xenopus pallium (l) and zebra finch HVC (o). (mouse, n = 3; Xenopus, n = 3; zebra finch, n = 6).

The initial infection of a target cell depends on interactions between viral envelope proteins and host-cell receptors. Because these host receptors vary across cell types and species, inefficient SINV transduction may result from limited compatibility between the viral structural proteins and the available cellular receptors. The MAPseq SINV genome can be packaged in, i.e., pseudotyped with, the structural proteins of other related alphaviruses^24–26^, a family of over 30 viruses that infect vertebrate and invertebrate hosts^27,28^. Pseudotyping leaves the MAPseq barcode expression and intracellular replication properties untouched while changing its tropism to that of the structural protein donating alphavirus^24–26^. It thus effectively uncouples MAPseq tracing from viral tropism.

We reasoned that if we pseudotyped the SINV MAPseq genome with the structural proteins of a range of different alphaviruses, we might be able to sidestep tropism issues and enable barcoding across a wide range of species (**Fig. 1b**). We initially selected five well-studied viruses—SINV itself, Venezuelan equine encephalitis viruses (VEEV), Western equine encephalitis viruses (WEEV), Eastern equine encephalitis viruses (EEEV) and Semliki Forest virus (SFV) —and used their structural proteins to package the propagation incompetent SINV MAPseq genome by swapping them into the DH-BB(5’Sin;TE12ORF) helper RNA (**Fig. 1c-d**)^23^. In what follows, we will refer to a specific pseudotyped virus by [structural protein donating virus]/[packaged genome], such that, e.g., VEEV/SINV refers to VEEV pseudotyped SINV virus. Each pseudotyped virus displayed distinct infection efficiencies in BHK-21 cells (**Supp. Fig. 1a**) and was produced at high titer as assessed by qPCR (**Fig. 1e**), demonstrating that structural proteins from diverse alphaviruses can efficiently package the MAPseq SINV genome. Note that all produced viruses are propagation incompetent and do not spread after initial infection (**Supp. Fig. 2a-d**). The viruses presented here thus do not pose biosafety concerns beyond standard Biosafety Level 2 containment.

**Figure 2.**
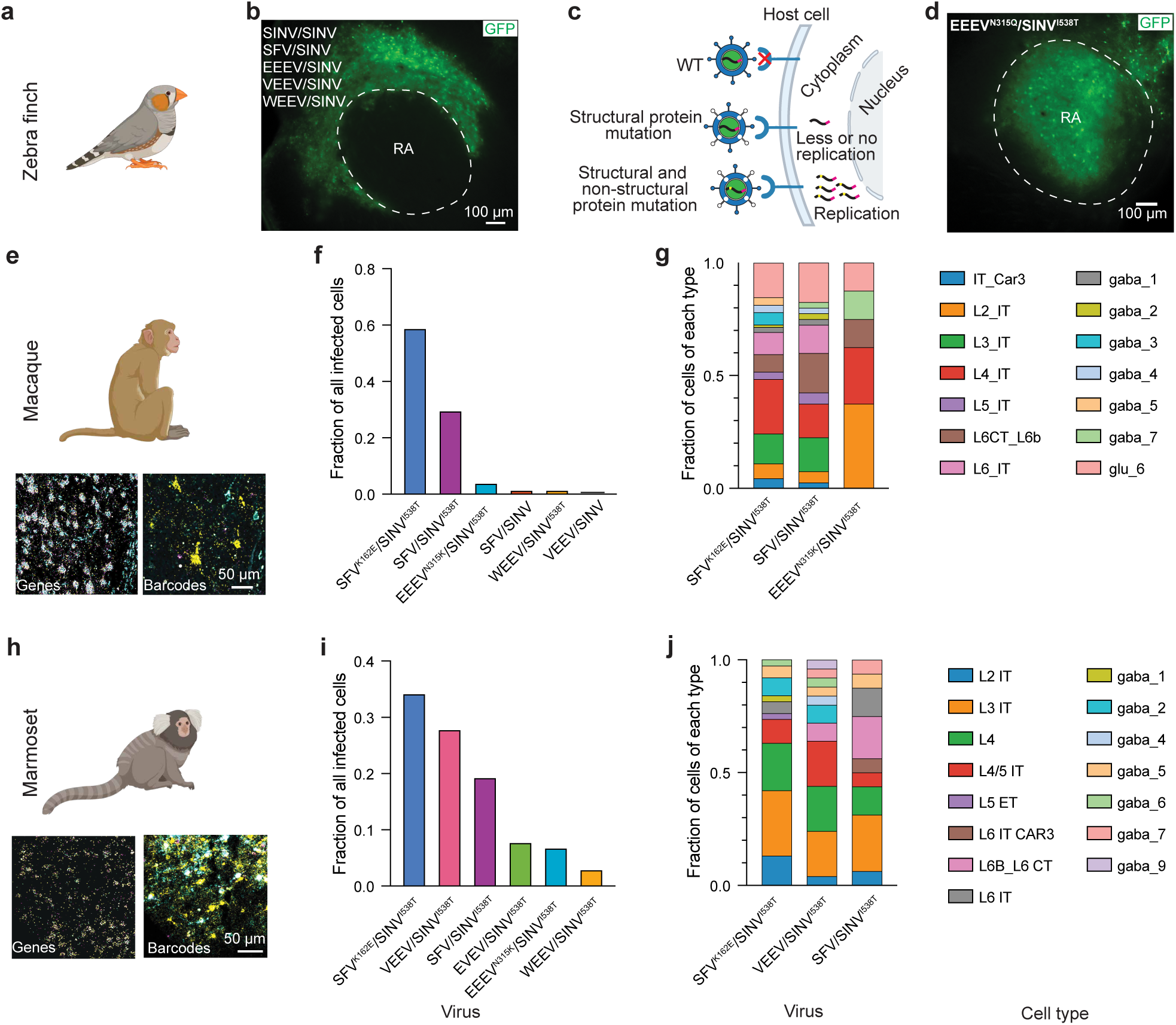
Pseudotyping and modulating the host response enables barcoding of a broader range of vertebrates. (a, b) Conventional pseudotyped viruses show no infection in zebra finch RA 24 h after infection. (c) Strategy for enhancing viral infectivity across species by introducing mutations into structural and nonstructural proteins. (d) EEEV^N315Q^/SINV^I538T^ produces robust infection in zebra finch RA 24 h after infection. (e-j) Screening of engineered viruses in macaque and marmoset frontal cortex. Representative *in situ* sequencing images showing viral barcodes and gene expression for macaque (e) and marmoset (h) after 72 h and 48 h of infection, respectively. SFV^K162E^/SINV^I538T^ produces robust infection in both species and broadly infects neuronal cell types (f, i).

To efficiently measure the tropism of these pseudotyped viruses in vivo while avoiding injection variability and minimizing the number of animals needed, we devised a pooled strategy that leverages the fact that individual SINV-based viral particles can drive the expression of a single barcode per infected target cell^17,24,29,30^. We produced each pseudotyped virus with a single, unique MAPseq identifier barcode, pooled barcoded viruses at equal titer, and delivered the mixture to the brain region of interest. After 24-72 hours of expression depending on transduced cell health, we then sequenced the identifier barcode and measured endogenous gene expression of each infected cell using BARseq^19,20^ *in situ* sequencing (**Fig. 1f**). In a single experiment, we therefore simultaneously quantify the infection efficiency and the cell-type tropism of all injected viruses. We first applied this strategy to mouse primary motor cortex (N=3; **Fig. 1g-i**), where we know that regular MAPseq virus (SINV/SINV) works well^24,31,32^. Indeed, SINV/SINV emerged as one of the more effective viruses alongside VEEV/SINV and EEEV/SINV in our screen (**Fig. 1h**). While VEEV/SINV infected more cells overall than SINV/SINV, it was not as specific to neurons as SINV/SINV (**Fig. 1h**), which had been experimentally evolved for increased neurovirulence in mouse cortex^33^. In a single injection, different viruses might compete with one another, potentially biasing pooled tropism measurements. However, the goal here is to identify a virus that efficiently infects target cells, rather than to establish an absolute rank of infectivity of viruses. Competition should thus not affect the end result. As expected, therefore, individual injections of VEEV/SINV and EEEV/SINV into motor cortex both yielded robust infection (**Fig. 1i**; **Supp. Fig. 1b**). Together, these results validate our viral screening approach.

To discover viruses that work well in other vertebrate clades, we then applied the same pooled strategy in the medial pallium of the amphibian *Xenopus tropicalis* (N=3) and the forebrain nucleus HVC of the zebra finch songbird *Taeniopygia guttata* (N=6). In both species we identified EEEV/SINV and SFV/SINV as the most efficient viruses (**Fig. 1j-l** and **m-o**). In contrast, regular SINV/SINV did not yield good infection, consistent with our previous observations (**Fig. 1a**). Independent single-virus validation experiments confirmed the pooled screening results (**Fig. 1l, o**; **Supp. Fig. 1c-e**). In each species and independent of the virus, only a single barcode was expressed per cell in the vast majority of cells (**Fig. 1g, j, m**), thus maintaining the one-barcode-per-cell infection regime exploited by MAPseq^24^.

Together, these findings establish a simple and generalizable strategy that combines alphavirus pseudotyping with pooled screening for identifying viruses suitable for neuronal barcode delivery across diverse vertebrate species.

### Viral mutations broaden the host range of MAPseq viruses across vertebrates

After viral entry, the virus needs to overcome the host innate immune defenses for expression and successful transduction^34,35^. While the host antiviral response did not restrict infectivity in the regions and species tested above, this response can be uneven across brain cell types even within the same species^34^. Indeed, when we injected different pseudotyped viruses into the mouse cerebellar nuclei, none of them yielded robust neuronal infection (**Supp. Fig. 3a**). Similarly, when we screened pseudotyped viruses in the zebra finch robust nucleus of the arcopallium (RA), a key nucleus in the zebra finch song circuit, we observed no transduction (**Fig. 2a-b**), even though we achieved good infection in the HVC of the same species (**Fig. 1o**; **Supp. Fig. 1d**). A primary driver of the host antiviral response is type I interferon signaling^36^. To test if failure of transduction was in fact due to this pathway, we first focused on mice where IFNAR1 knockout animals with a disabled interferon type 1 response are available^37^. In contrast to our results in wild type animals (**Supp. Fig. 3a**), injection of EEEV/SINV, but not SINV/SINV (**Supp. Fig. 3b, c**), into the cerebellar nuclei of these knockout animals resulted in robust infection of cerebellar nuclei neurons. Variation in interferon signaling across species and brain regions therefore acts as a second determinant of MAPseq viral tropism.

**Figure 3.**
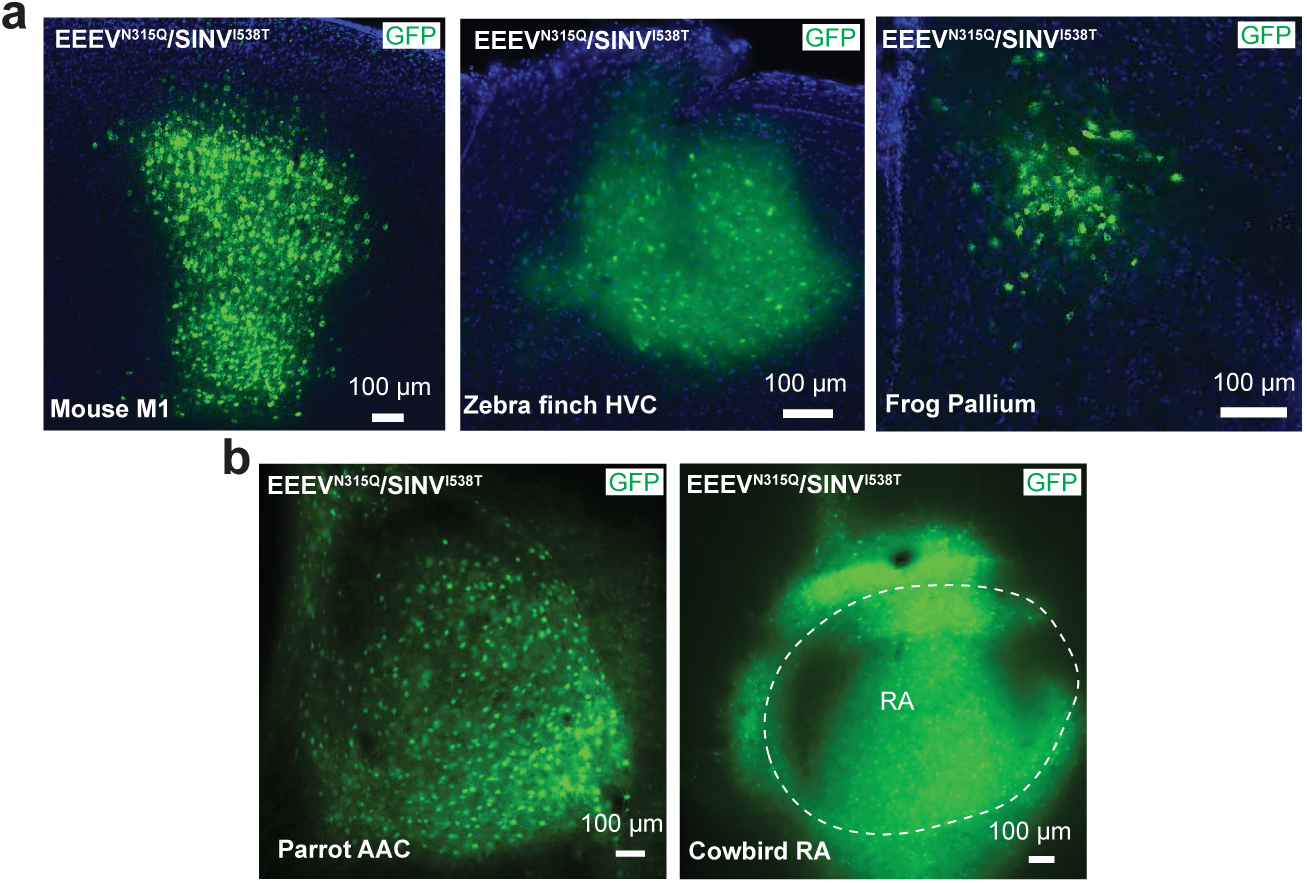
A single engineered virus enables infection of highly divergent species. (a, b) EEEV^N315Q^/SINV^I538T^ produces robust infection across diverse vertebrate species and brain regions, including mouse M1, zebra finch b HVC, and Xenopus pallium (a), as well as the previously untested budgerigar AAC and cowbird RA (b).

Genetically disrupting host interferon signaling to facilitate viral infection is not an option in most species, including zebra finches. However, different point mutations in the SINV nonstructural proteins have been identified that modulate its interaction with the host, potentially providing a way to overcome antiviral responses by changing the virus rather than the host^38^. In particular, it has been reported that changing amino acid 538 from threonine (T) to isoleucine (I) in the AR86 SINV strain reduces viral expression in interferon-competent cell lines^39^. Amino acid 538 is an I in the TE12 Sindbis strain used in MAPseq^23^, suggesting that reverting this position to a T might enhance the virus’s ability to overcome host antiviral responses. Indeed, EEEV/SINV^I538T^ produced robust neuronal infection in the cerebellar nuclei, even in wild type mice (**Supp. Fig. 3d**), demonstrating that this single point mutation in the MAPseq genome can overcome interferon-imposed tropism barriers and obviate the need for IFNAR1 knockout animals.

To further increase our chances of finding an infective pseudotyped virus in hard-to-infect brain regions, we also increased our pool of structural proteins for pseudotyping. We included Everglades virus (EVEV), an alphavirus that has been reported to infect birds^40^, and modified SINV, SFV, and EEEV E2 structural proteins at sites expected to yield increased cell entry. Specifically, we included SINV^N196Q, N318Q^ and SFV^K162E^ mutants, which were shown to increase virulence^41,42^. Additionally, we included EEEV^N315Q^ and EEEV^N315K^ mutants, motivated by the observation that the EEEV E2 glycan at N315 is structurally buried, unlike those of other alphaviruses^43^.

Combining the extended pseudotyping panel and improved defense against interferon signaling, we produced barcoded pseudotyped viruses using the SINV^I538T^ genome and packaged them in wild type and mutant structural proteins (**Fig. 2c**; **Supp.** **Fig. 4a**-**b**). We then injected these pseudotyped viruses into the previously uninfectable zebra finch RA. By screening for GFP expression after injecting different combinations of viruses, we found that EEEV^N315Q^/SINV^I538T^ uniquely and robustly infected the previously recalcitrant RA cells (**Fig. 2d**; **Supp. Fig. 4c-f**) showing that both increased viral entry and defense against host repones were required to infect this brain region.

**Figure 4.**
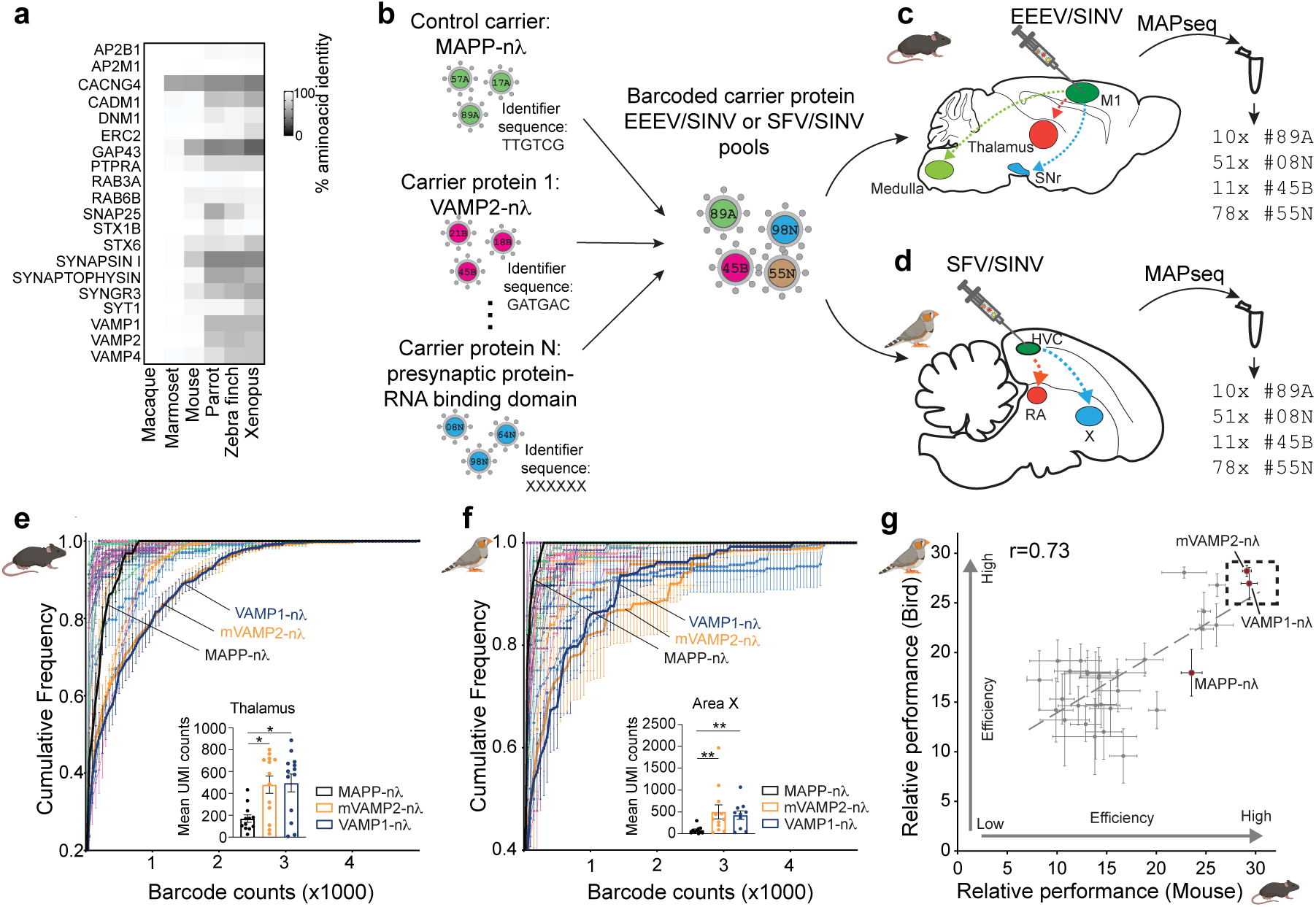
VAMP-based carrier proteins work efficiently across distantly related species. (a) The presynaptic proteins used as basis for carrier proteins are highly conserved across vertebrates. (b-d) Design of the pooled carrier protein screen in mouse motor cortex and zebra finch HVC. (e, f) Cumulative distribution of barcode UMI counts in target regions for each carrier protein, with inset showing mean barcode counts for the MAPP-nλ control and top performers for mice (e) and zebra finches (f). VAMP1-nλ and mVAMP2-nλ show substantially improved performance relative to MAPP-nλ in both species. (Mouse: n = 13 hemispheres, Zebra finch: n = 12 hemispheres, Mann-Whitney test, *p<0.05, **p<0.01) (g) Scatter plot of the average ranking of each carrier protein across target regions in each species, showing high correlation of ranks across mice and zebra finches and consistent top performance of VAMP1-nλ and mVAMP2-nλ.

We next asked whether this improved viral pool could also yield viruses that effectively transduce neurons in non-human primates, where initial MAPseq experiments with the original SINV/SINV constructs showed relatively low transduction efficiencies^22^. We screened a combined pool of wild type and SINV^I538T^ genome pseudotyped viruses in macaque frontal cortex and a SINV^I538T^ virus pool in marmoset frontal cortex (**Fig. 2e-j**; **Supp.** **Fig. 4g**-**h**). In both primate species, SFV^K162E^/SINV^I538T^ was the top performer and infected a broad range of neuronal cell types, vastly outperforming SINV/SINV.

Taken together, our results demonstrate that by modifying the structural proteins and enhancing the anti-host response of the Sindbis genome we can infect previously difficult to infect brain regions and species and identify at least one virus to efficiently barcode neurons in every brain region and vertebrate animal we tested.

### A single virus can infect species separated by 350 million years of evolution

To simplify circuit tracing in diverse species, it would be advantageous to identify a single virus that overall is most general in its infectivity, as a “first best guess” for what pseudotyped virus to try in a previously untested brain region or species without the need for screening. We therefore tested EEEV^N315Q^/SINV^I538T^, the virus that infected the hard-to-infect bird RA and mouse cerebellar nuclei, for infection in mouse M1, zebra finch HVC, and Xenopus pallium. We also tested it in previously untested species, namely in the budgerigar parrot (*Melopsittacus undulatus*) central nucleus of the anterior arcopallium (AAC) and the cowbird (*Molothrus ater*) RA. Remarkably, in all taxa (mice, birds, frogs), species, and regions EEEV^N315Q^/SINV^I538T^ led to robust infection (**Fig. 3a-b**; **2d**). Together with SFV^K162E^/SINV^I538T^ which efficiently infected the two non-human primate species we tested, these results suggest that one or a few virus pseudotypes may be sufficient to achieve infection across a broad range of vertebrate species and brain regions.

### Identification of an effective carrier protein across species

Beyond infection, single-neuron projection mapping by barcoded connectomics requires efficient trafficking of barcode mRNA into axon terminals. The original MAPseq method used MAPP-nλ as a carrier protein to traffic barcode mRNAs into axons^17^. MAPP-nλ is a chimeric protein based on the pre-mGRASP^44^ presynaptic protein fused to the RNA binding domain nλ, which binds the boxB hairpin on the barcode mRNA and moves it into axonal terminals. However, whether this protein works well in carrying barcodes into axon terminals in diverse species is unknown.

To identify a carrier protein that works efficiently across species, we performed a barcoded cell autonomous screen in mice and zebra finches. These two species are separated by over 300 million years of evolution, and we reasoned that any carrier proteins that performed well in both systems would likely be effective carrier proteins in a wide variety of regions and species. As a basis for potential carrier proteins, we selected 20 presynaptically localized proteins that are highly conserved across vertebrate species (**Fig. 4a**). A protein from one species would therefore likely traffic similarly in another, biasing our screen towards carriers that perform well across species. We fused the macaque sequence of each protein to one or four copies of RNA binding domain nλ, or to one copy of RNA binding domain PP7 (**Supp. Fig. 5a**), with most constructs using one copy of nλ (**Supp. Fig. 5d**). Finally, we inserted them into a MAPseq genome in place of MAPP-nλ with matching binding sites in the barcode mRNA. We also included the original MAPP-nλ and the mouse VAMP2 (mVAMP2)-nλ carrier used in a recent axonal BARseq study^45^ as internal controls. This yielded a total of 31 potential carrier protein constructs. We produced plasmid barcode libraries for each construct individually with a minimum diversity of 5×10^6^ (**Supp. Fig. 5d**). Each random barcode ended in a unique known identifier sequence (**Supp. Fig. 5a, d**) that reported the carrier protein used in each construct. We then generated individual viral libraries for each construct using both EEEV/SINV and SFV/SINV, two pseudotypes that showed efficient infection in mouse and zebra finch (**Supp. Fig. 1b, d**). For each pseudotype, we pooled the viruses at matched titers to ensure an even representation of all carriers in the final viral pool.

**Figure 5.**
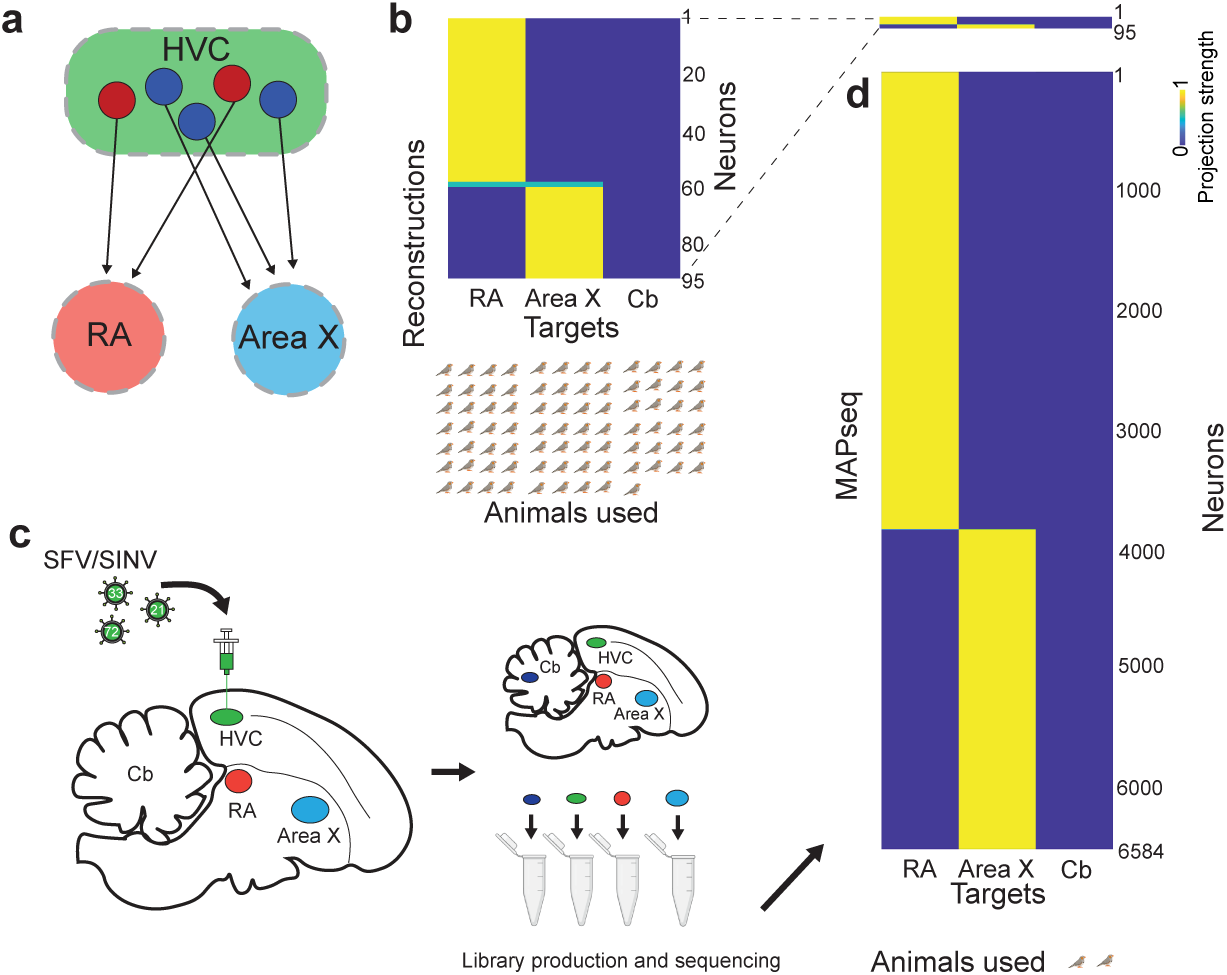
Universal MAPseq reagents enable efficient single-neuron projection mapping from zebra finch HVC. (a) Schematic of the HVC neuronal projection pattern that is dominated by dedicated projections to RA and Area X. (b) Binary single-neuron projection matrix produced from ground-truth single-neuron reconstructions (N = 95 neurons) in 81 birds, based on data collected by Benezra et al., 2018. Almost all HVC neurons project to either RA or Area X, with only rare neurons (2/95) collateralizing to both. (c) Schematic of the experimental workflow for mapping neuronal projections from zebra finch HVC using MAPseq reagents developed above (d) Single neuron projection matrix produced by MAPseq showing that HVC neurons project to either RA or Area X, with only rare neurons projecting to both and no detectable projections to the negative control cerebellum (N = 6584 neurons, n = 3 hemispheres, 2 birds). These results match well to the ground truth data, but required much fewer animals to obtain

We then injected the EEEV/SINV and SFV/SINV virus pools into mouse primary motor cortex (N=13 hemispheres) and zebra finch HVC (N=12 hemispheres), respectively. After viral expression, we dissected the distal motor cortex target regions thalamus, substantia nigra pars reticulata (SNr), and medulla in the mouse, and HVC targets Area X and RA in the zebra finch, as well as the injection site and sequenced the barcodes contained therein using the MAPseq2 protocol^32^ (**Fig. 4b-d**; **Supp. Fig. 5b, c**). We then assigned each barcode to its corresponding carrier protein based on the identifier sequence and quantified carrier protein performance by the cumulative distribution of molecule counts per barcode in each target region. Different carrier proteins showed different abilities to carry barcode mRNA into target regions, with some under-and others outperforming the MAPP-nλ control (**Fig. 4e-f**; **Supp. Fig. 5e-i**). The performance ranking of carrier proteins based on the average UMI count of barcodes carried by them was highly reproducible across target brain regions in the same species (**Supp. Fig. 6a-b**). Remarkably, different carrier proteins also performed similarly across mice and zebra finches (**Fig. 4g**; **Supp. Fig. 7a-e**), likely reflecting the deep conservation of presynaptic protein trafficking in vertebrates and indicating that the same carrier proteins will work well across a broad range of animals. In particular, VAMP1-nλ and mVAMP2-nλ were consistently among the most efficient carrier proteins in both mice and zebra finches across regions (**Fig. 4g**; **Supp. Fig. 6a-b and 7a-e**), carrying significantly more barcode molecules into target regions than MAPP-nλ (**Fig. 4e,f**; **Supp. Fig. 5f, g, i**). Packaged in any of the effective viral pseudotypes identified above, these two carrier proteins should therefore support barcoded connectivity mapping in a broad range of species.

### Mapping single-neuron projections in birds

To validate our combined reagents, we sought to map the single-cell projections of zebra finch HVC. HVC is a brain region central to song learning and production in zebra finches^46,47^, and one of only a few brain regions in non-mammalian vertebrates for which strong single-cell projection ground truth data exist: individual HVC neurons project to either Area X or RA, and almost never both(**Fig. 5a**)^48,49^. In particular, Benezra and colleagues reconstructed the projections of 95 HVC neurons in 81 birds, yielding a gold standard comparison dataset to our MAPseq tracing (**Fig. 5b**)^49^. To compare to this, we injected SFV/SINV barcoded MAPseq library containing the mVAMP2-nλ carrier protein into HVC, and dissected and sequenced the barcodes found in target regions Area X and RA, as well as the cerebellum as a negative control after 24 hours of infection (**Fig. 5c**). Based on these sequencing data, we then quantitatively reconstructed each barcoded neuron’s projection pattern. In total, we mapped the projections of 6,584 HVC neurons across three hemispheres of two birds. In each hemisphere, almost all MAPseq neurons projected either to Area X or RA (**Fig. 5c-d**; **Supp. Fig. 8a-c**), as expected. In addition to these classic, dedicated projection neurons, we also recovered a small number of neurons (<1%) that bifurcate and project to both RA and X (**Supp. Fig. 8d-f)**, matching the Benezra dataset, that includes two such non-classical neurons. Our MAPseq results thus replicate the ground truth data collected by traditional, gold standard single neuron reconstruction methods, demonstrating that our new MAPseq reagents can robustly and accurately map neuronal projection in birds. At the same time, this first barcoding-based single-neuron projection dataset from a non-mammalian vertebrate already provides a ∼70× increase in traced HVC neurons available in the literature while using 40× fewer animals, highlighting the advantages of barcoded connectivity tracing.

## Discussion

Here, we overcome a central bottleneck in neuroscience investigations by developing a broadly applicable, ready-to-use tool for mapping single-cell projection patterns at scale for thousands of neurons per animal in any laboratory in a range of vertebrates from frogs to birds to nonhuman primates. We establish a general solution to viral tropism by viral pseudotyping and modulating the anti-viral host response and identify a set of carrier proteins that are effective across highly divergent species. Together, we provide a universal, modular MAPseq platform in which a common barcoded genome encoding a broadly effective carrier protein is packaged with one or two sets of engineered structural proteins to enable out-of-the-box single-cell connectivity mapping across many brain regions in species separated by over 350 million years of evolution.

While recent work has expanded the range of species AAV can be used in^50–52^, single-neuron projection mapping in species other than mice is still very rare. This is because AAV-based modern fluorescence-based single-cell tracing tools require significant optimization in each species, large investments in infrastructure, and remain difficult to scale and perform in individual labs^53,54^. Compounding this issue, AAV depends not only on efficient transduction but also on species-compatible promoters to drive transgene expression – an especially challenging requirement in non-traditional organisms where promoter activity is often poorly characterized^50,55^. In contrast to these traditional neuroanatomical tools, barcoded connectomics is an attractive candidate for single-neuron projection mapping in diverse species, as it readily and rapidly provides the scale and single-cell resolution needed to answer biological questions in any laboratory. Moreover, barcode and carrier protein expression are driven by the virus’ own strong promoter, dispensing with the need for species-specific promoters. By decoupling the viral delivery machinery from the barcoded payload into a modular system and identifying a general carrier protein, our work now enables barcoded projection mapping across multiple regions in diverse species. As no modifications to the barcode expressing payload are required in different species, data collected in different animals is fundamentally comparable, a critical feature for single-cell comparative connectomics.

Our work in extending MAPseq barcoding across vertebrates also immediately extends the reach of other barcode-based technologies to these species. In particular, *in situ* sequencing methods BARseq2^19^ and BARseq3^20^, and single-cell sequencing method ConnectID^56^ have been developed to simultaneously measure a cell’s gene expression and its barcoded single-cell projection pattern for a comprehensive understanding of cell type – a particularly valuable combination of readouts in species without extensive collections of cell-type-specific reagents. Their application so far, however, has been limited to mice, due to the difficulty of barcoding other species. This limitation is now removed by the work presented here, already enabling simultaneous projection and gene expression mapping of single cells in marmoset thalamus in a concurrent study^57^.

Finally, while we present barcoding data for seven highly divergent vertebrate species, we expect our modular system to be readily adapted to any other vertebrate species of interest. For viral infection in a new species, EEEV^N315Q^/SINV^I538T^ pseudotyped virus should serve as a first “best-guess” virus, as it performed robustly in all brain regions and species we tested. In scenarios where it underperforms, the 20 viral variants screened in this study can serve as a practical starter set for screening in new species. For barcode trafficking, our results indicate that VAMP1-nλ and mVAMP2-nλ work efficiently as carrier proteins across highly divergent animals. This likely holds true for any vertebrate, as VAMP family proteins, part of the SNARE complex of any synapse, are highly conserved across species^58,59^. We therefore expect that one or a small handful of pseudotyped viruses carrying a VAMP1/2-nλ expressing barcoded genome will be sufficient to enable barcoded connectomics across all vertebrates, paving the way for comparative neuroanatomy at single-neuron resolution.

## Materials and Methods

### Animal husbandry

All animal procedures were approved by the University Animal Care and Use Committees at Johns Hopkins University, Mount Sinai, NIH, or New York University, and were performed in accordance with NIH standards and the approved protocols for mice, zebra finches, budgerigars, cowbirds, Xenopus, macaques, and marmosets. C57BL/6J (JAX #000664) and INFARI KO (JAX #028288) mice were purchased from Jackson Laboratory. Adult male zebra finches and budgerigars were purchased from Maryland Exotic Birds and adult female Xenopus tropicalis were purchased from Xenopus1. Cowbirds were captured in the wild in Pennsylvania per our protocol at New York University. The macaque was purchased from the National Biomedical Research Institute at the University of California, Davis. The marmoset was from the NIH internal breeding colony. All the animals were maintained at a 12-h light-dark cycle with food and water in species appropriate conditions.

### Plasmids

#### For Virus screening

We generated helper constructs for pseudotyping based on the DH-BB(5’Sin;TE12ORF) construct^23^ (Addgene # 72309) by replacing the structural protein open reading frame with that of other alphaviruses. Structural protein sequences were obtained from NCBI (EVEV: NP_040824.1; EEEV: NP_632022.1; VEEV: NP_040824.1; WEEV: NP_640331.1; SFV: NP_463458.1) and synthesized by Twist Biosciences. We generated mutant structural protein DNA fragments of SFV (K162E), SINV (N196Q, N318Q, N196K, N318K), and EEEV (N315Q, N315K) by overlap PCR and inserted fragments between NcoI (New England Biolabs, R0193S) and EcoRV (New England Biolabs, R3195S), BlpI (New England Biolabs, R0585S) and BssHII (New England Biolabs, R0199S), and BbvCI (New England Biolabs, R0601S) and BsiWI (New England Biolabs, R3553S) restriction sites, respectively. We generated I538T mutant non-structural protein by overlap PCR and inserted the fragment between BbvCI and ClaI (New England Biolabs, R0197S). We inserted a unique barcode sequence (Supplemental Table 1) between NotI (New England Biolabs, R3189L) and MluI (New England Biolabs, R3198L) restriction sites in the HZ120 (Addgene # 265231), SINV and HQ065 (Addgene # 262100) SINV^I538T^ MAPseq genomic construct for each virus.

### For Carrier protein screening

We ordered the carrier protein sequences as DNA fragments from Twist Bioscience and cloned each fragment into the pSinEGdsp construct jk100 (Addgene # 79785)^17^ to replace original MAPP-nλ carrier protein. To generate the carrier protein sequences, we fused the RNA-binding protein domain nλ or PP7 to either the N-or C-terminus of each synaptic protein at positions previously reported to tolerate the insertion of fluorescent proteins. Detailed information on the fusion sites is provided in the corresponding Addgene plasmid sequence and Supplemental Table 3. The NCBI accession numbers for the synaptic proteins used are listed in Supplemental Table 3. For each construct, we then produced high diversity plasmid libraries by inserting a diverse pool of random DNA oligos, as described previously^17,60^. The barcode was designed as was N20-X6, where X are bases specifying the identifier tag used to denote each individual carrier protein (Supplemental Table 1). Briefly, we synthesized single-stranded DNA oligos as ultramers from IDT. We then generated double strand DNA by 15 cycles of PCR amplification using Accuprime Pfx (Invitrogen, 123444024) and only the reverse primer (5’-GGT AGC GGC CGC AGT AC-3’), and then removed single stranded DNA with Thermolabile Exonuclease I (New England Biolabs, M0568S). We purified the product using SV Wizard PCR cleanup kit (Promega, A9282). We then digested plasmid backbone and the dsDNA fragments by NotI and MluI at 37 °C overnight and purified the reaction products using SV Wizard PCR cleanup kit. We performed ligation for inserts and backbone at ratio around 3-5 using T4 ligase (New England Biolabs, M0202M) at 16 °C overnight. We cleaned up ligation reactions using AMPure XP beads (Beckman Coulter, A63881) and digested the products by Plasmid Safe ATP Dependent DNase (Biosearch Technologies, E3101K). We cleaned up the products again using AMPure XP beads. We performed electroporation of MegaX electrocompetent cells (Thermo Fisher, C640003) and DNA ligation product in a 0.1 cm cuvette (Bio-Rad, 1652089) following the manual. After electroporation, we immediately added SOC medium and incubated bacteria in LB medium containing carbenicillin at 37 °C for 12 hours with shaking at 200 rpm.

### Virus production

We used BHK cells to produce pseudotyped MAPseq viruses, as previously described^17^. Briefly, helper plasmids and genome plasmids were linearized by XhoI (New England Biolabs, R0146L) and PacI (New England Biolabs, R0547L) respectively and helper and genomic RNA were produced by in vitro transcription using the mMessage Machine SP6 kit (Invitrogen, AM1340) according to the manufacturer’s instructions. We cleaned the RNAs with AMPure XP beads (Beckman Coulter, A63881) and transfected the RNA into BHK cell with a 1.7:1 weight ratio of genomic to helper RNA using Lipofectamine 2000 (Invitrogen, 11668027). We collected supernatant 40 hours after transfection and concentrated the virus by ultracentrifugation at 160,000 × g, at 4 °C for 2 hours. We removed the supernatant, leaving approximately 40 µL, and then placed the tube on ice for 3 hours to resuspend the virus. We aliquoted each virus into small volumes and stored them at -80 °C.

Virus titer was measured by qPCR as previously described^17^. Briefly, we treated virus with RNase If I (New England Biolabs, M0243S) at 37 °C for 30 min and extracted protected genomic RNA using Trizol. We performed reverse transcription using RT primer 5’-GGG TCG CCT TGC TTG AAG-3’ and SuperScript™ IV Reverse Transcriptase (Thermo Fisher, 18090050) following the manufacturer’s instructions. We performed SYBR green qPCR using the reverse transcription products with Forward primer 5’-TAT CCG CAG TGC GGT TCC AT-3’ and Reverse primer 5’-TGT CGC TGA GTC CAG TGT TGG-3’ and compared these resulting Ct values to serial dilutions of the corresponding genomic plasmid to calculate an absolute viral titer in genome copies/µL.

### Plaque assay

We performed plaque assays as previously described^23,24^. Briefly, we infected BHK-21 cells at approximately 90% confluence with serial dilutions of SINV, SFV^K162E^/SINV^I538T,^ EEEV/SINV, or EEEV^N315Q^/SINV^I538T^ to ensure well-separated plaques in 24-well plates. After 1 hour infection, we overlaid cells with 0.4% low melting point agarose (Invitrogen, 16520050) prepared in cell culture medium [10% FBS (Millipore Sigma, F4135) in DMEM (Thermo Fisher, 11965092)]. After 24 hours, we imaged 3three fields of view.

### Surgery and injections

We mixed equal genome copies of each virus for virus screening and carrier protein screening before injection.

### Mice

We anesthetized 8-to 10-week-old male C57BL/6J mice with isoflurane (2.5% for induction and 1.5% for maintenance). We then injected 400 nL of the pseudotyped virus pool (SINV/SINV, EEEV/SINV, VEEV/SINV, WEEV/SINV and SFV/SINV), EEEV/SINV carrier protein virus pool, or a single pseudotyped virus into the primary motor cortex (AP: +1.0 mm, ML: ±1.6 mm, DV: -1.0 mm) for virus screening, carrier protein screening, or individual tests, respectively. We injected individual viruses (SINV/SINV, EEEV/SINV, VEEV/SINV, WEEV/SINV, SFV/SINV, or EEEV/SINV^I538T^) into IFNAR1 KO and wild type mouse cerebellar nuclei (relative to lambda, anterior medial nucleus: AP: -1.83 mm, ML: ±0.75 mm, DV: -3.4 mm; posterior medial nucleus: AP: -2.18 mm, ML: ±1.0 mm, DV: -3.2 mm; interposed nucleus: AP: -1.83 mm, ML: ±1.5 mm, DV: -3.3 mm; lateral nucleus: AP: -1.4 mm, ML: ±2.45 mm, DV: -3.8 mm) to test role of the host’s antiviral response in limiting infection.

### Zebra finches

We anesthetized adult male zebra finches with isoflurane (3.5% for induction and 2.2% for maintenance). We then injected 400 nL of the pseudotyped virus pool (SINV/SINV, EEEV/SINV, VEEV/SINV, WEEV/SINV and SFV/SINV), the SFV/SINV carrier protein virus pool, or an individual virus into HVC (beak angle 45°, AP: 0 mm, ML: ±2.2 mm, DV: -0.3 mm) for virus screening, carrier protein screening, or individual tests, respectively. We injected 500nL of EEEV^N315Q^/SINV^I538T^ into RA using the following coordinates: beak angle 45°, AP: -0.6 mm, ML: ±2.35 mm, DV: -2.78 mm. For MAPseq projection mapping, we injected 500 nL of an SFV/SINV barcoded virus library into HVC.

### Cowbirds

We anesthetized adult male cowbirds using 1-3% isoflurane in oxygen. Animals were positioned in a stereotaxic apparatus with the beak bar set at an angle of 35°. At this angle, RA was targeted using the following coordinates: AP, -0.25 mm; ML, ±2.75 mm; and DV, -2.7 to -3.5mm. After the pipette was lowered into RA, 500 nL EEEV^N315Q^/SINV^I538T^ barcoded virus library was injected.

### Budgerigars

We targeted AAC in budgerigars following our previously published procedures^61^. Briefly, adult male budgerigars were anesthetized with 1-3% isoflurane. We then first localized AAC using a silicon probe at the following coordinates: ML, 5 to 6 mm; AP, 5 to 6 mm; DV, -3.2 to -3.8 mm. After AAC was identified electrophysiologically, an injection pipette was lowered into the target, and 500 nL EEEV^N315Q^/SINV^I538T^ barcoded virus library was injected.

### Xenopus

We deep anesthetized adult female *Xenopus tropicalis* by immersion in MS222 (1 g/L) solution for 10 minutes. The MS222 was prepared with dechlorinated water and buffered to a pH of 7.0 with sodium bicarbonate. We injected 400 nL of the pseudotyped virus pool (SINV/SINV, EEEV/SINV, VEEV/SINV, WEEV/SINV and SFV/SINV), or individual viruses into the pallium (AP: +4.5 mm, ML: ±0.3 mm, DV: -0.7 mm).

### Marmosets

We anesthetized a 16-month-old male marmoset with alfaxalone (8 mg/kg) and diazepam (1-3 mg/kg) and maintained with 0.5-2% inspired isoflurane. We administered a broad-spectrum antibiotic (Norocillin; Norbrook, Lenexa, KS, USA; 27 mg/kg, 0.1 mL, i.m.), atropine (20 μg/kg, i.m.), and dexamethasone (0.3 mg/kg, i.m.). We then secured the animal in a stereotaxic frame and continuously monitored its physiological status, including oxygen saturation (SpO₂) and core body temperature, maintaining these parameters within predefined limits. We made a small craniotomy and injected 500 nL of the pseudotyped virus pool (SINV/SINV^I538T^, EEEV/SINV^I538T^, WEEV/SINV^I538T^, VEEV/SINV^I538T^, SFV/SINV^I538T^, EEEV^N315Q^/SINV^I538T^, _SINV_N196Q;N318Q_/SINV_I538T_, SFV_K162E_/SINV_I538T_, SINV_N196K;N318K_/SINV_I538T_, EEEV_N315K_/SINV_I538T_, and_ EVEV/SINV^I538T^) into the frontal cortex with a rate of 180 nL/min at a depth of 1 mm below the brain surface using a 5 μL Hamilton syringe with a 15 degree beveled 33 gauge needle. After injection, we sealed the craniotomy with Kwik-Sil and closed the scalp incision with intradermal sutures (coated Vicryl 5-0). We then allowed the animal to recover in a temperature- and humidity-controlled cage.

### Macaques

We anesthetized a 9-year-old male macaque (*Macaca mulatta*, 9.6 kg) with ketamine (10 mg/kg) and dexmedetomidine (0.015 mg/kg) and maintained the plane of anesthesia with 1-3% isoflurane. During an aseptic neurosurgery, the skin and fascia were opened and the muscles were retracted. A small craniotomy was made over the principal sulcus and injected the pseudotyped virus pool (SINV/SINV, EEEV/SINV, VEEV/SINV, WEEV/SINV, SFV/SINV, EEEV^N315Q^/SINV, SINV^N196Q;N318Q^/SINV, SFV^K162E^/SINV, EVEV/SINV, SINV/SINV^I538T^, EEEV/SINV^I538T^, WEEV/SINV^I538T^, VEEV/SINV^I538T^, SFV/SINV^I538T^, EEEV^N315Q^/SINV^I538T^, _SINV_N196Q;N318Q_/SINV_I538T_, SFV_K162E_/SINV_I538T_, SINV_N196K;N318K_/SINV_I538T_, EEEV_N315K_/SINV_I538T_, and_ EVEV/SINV^I538T^) was injected at two locations into the left frontal cortex using a 30 gauge Hamilton syringe with a 45 degree bevel. We performed injections at two depths, 2 mm and 1 mm below the cortical surface, with 1.5 μL of virus delivered at each depth at a rate of 0.5 μl/min. At the completion of injections, the muscles, fascia and skin were sutured, the animal recovered and returned to their home cage where they were closely monitored until normal behavior resumed. Postoperative medications were provided including buprenorphine (0.01 mg/kg, i.m., every 8 hours) and meloxicam (0.2 mg/kg, i.m., every 24 hours), based on attending veterinary guidance, as well as cefazolin (25 mg/kg, i.m., every 24 h) and dexamethasone sodium phosphate (0.4-1 mg/kg, every 12-24 hours) on a descending dose schedule.

For mouse, zebra finch, and Xenopus virus screening, we dissected brains 24 hours after infection and punched out the injection site. We then immediately embedded the tissue samples in O.C.T (Thermo Fisher, 23-730-571) and stored them into -80 °C. Marmoset virus screening was performed 48 hours post infection. Anesthesia was induced with an overdose of pentobarbital (>100 mg/kg). The animal was then transcardially perfused with ice-cold artificial cerebrospinal fluid (aCSF). The brain was extracted, dissected into blocks and flash-frozen in dry-ice cooled isopentane. Macaque virus screening was performed 72 hours post infection. Anesthesia of the macaque subject was induced with ketamine /dexmedetomidine (10 mg/kg and 0.125 mg/kg, respectively) and then sodium pentobarbital (100 mg/kg) was administered. The animal was then transcardially perfused with RNAse-free ice-cold PBS, after which the brain was extracted, dissected into blocks, and flash frozen in 2-methylbutane^22^. All brain blocks were stored at -80°C until processing.

For carrier protein screening in mouse and zebra finch, we dissected brains after 48 hours and 24 hours of infection, respectively. We then immediately embedded brains in O.C.T, flash-frozen them, and stored them at -80 °C.

For the testing of individual viruses in mice, Xenopus, zebra finch, budgerigar, and cowbird, we transcardially perfused the animals with PBS followed by 4% paraformaldehyde (PFA) (Booster Bio, AR1068). The brain was then cryoprotected in sucrose, frozen, and stored at -80 °C.

For MAPseq projection mapping, we dissected zebra finch brains after 24 hours of infection. We then embedded the brains into O.C.T, flash-frozen them, and stored them at -80 °C.

### Virus screening

#### BARseq3 library preparation and *in situ* sequencing

We performed viral screening in mice, zebra finches, and Xenopus using BARseq3^20^. Briefly, we designed the BARseq3 SNAIL probe hybridization sequences using Picky^62^ v2.2 and custom scripts as previously described^20^ with the following parameters. length: 40-46 nt; matched length: 10-15 nt; GC content: 30-70; minimum trigger similarity: 75; minimum temp difference: 15; DNA concentration: 1 nM; salt concentration: 500 mM. We generated SNAIL probes containing gene-specific ID sequence with a minimum Hamming distance > 2 between genes. We designed four probes for each gene (Mouse: Gad1, Slc17a7, Gfap, Tmem119; Zebra finch: Slc17a7, Slc17a6, Gad1, Gad2, Plp1, Sox9, Csf1r; Xenopus: Slc17a6, Gad1, Gad2, Snap25, Unc93b1, Slc4a4, Slc1a2, Cd74, Acsbg1, Slc32a1) and ordered probes as custom oPools oligo pools from IDT. All genes probed for and associated probe sequences can be found in Supplemental Table 2.

We then cryosectioned the samples at 20 µm thickness and hybridized probes [50 nM per oligo, 2× SSC (Sigma-Aldrich, S6639), 10% (vol/vol) formamide (Calbiochem OmniPur, 75-12-7), 10% (vol/vol) Triton X-100 (Sigma-Aldrich, 93443), 20 mM ribonucleoside vanadyl complex (New England Biolabs, S1402S), 0.1 mg/ml Salmon Sperm DNA (Invitrogen, AM9680), 0.2 U/μL Superasin RNase Inhibitor (Invitrogen, AM2696)] at 40 °C for 19-20 hours. We then washed the samples [4× SSC, 0.2 U/μL Superasin RNase Inhibitor, 1% Tween-20 (Millipore Sigma, P9416) in PBS] at 37 °C for 20 min, performed ligation [1× T4 DNA ligase buffer, 0.1 mg/ml recombinant albumin (New England Biolabs, B9200S), 0.2 U/μL Superasin RNase Inhibitor (Invitrogen, AM2696), 2 U/μL T4 DNA ligase (Thermo Fisher Scientific, EL0011)] at room temperature for 5 hours, followed by a first round of rolling-circle amplification (RCA) [1× Phi29 buffer (McLab, PP-200), 0.2 mg/mL BSA (New England Biolabs, B9200S), 0.4 mM dNTPs (Thermo Fisher Scientific, R0192), 250 μM aminoally-dUTP (Invitrogen, AM8439), 1 U/μL Phi29 polymerase (MCLab, PP-200), 0.2 U/μL Superasin RNase Inhibitor (Invitrogen, AM2696), 5% (vol/vol) Glycerol (Millipore Sigma,65516)] at 25 °C overnight. After crosslinking [50 mM BS(PEG)9 (Broadpharm, BP-21504), 20 mM RVC (New England Biolabs, S1402S), 0.5% Tween-20 in PBS] at room temperature for 1 hour, we quenched the samples with 1 M Tris-HCl for 30 min. We performed reverse transcription using the barcode specific primer (1× QuantumScript™ Reverse Transcriptase buffer, 0.2 mg/mL BSA, 1 U/μL RiboLock RNase inhibitor, 0.5 mM dNTPs, 1 μM barcode LNA primer, 20 U/μL QuantumScript™ Reverse Transcriptase) at 37 °C for 7 hours and then crosslinked the sample again with BS(PEG)9. We hybridized the barcode padlock probe followed by gap-filling and circularizing [1× Ampligase buffer, 50 mM KCl, 0.1 μM barcode padlock, 0.05 mM dNTPs, 5% (vol/vol) Glycerol, 20% (vol/vol) formamide, 0.5 U/μL Ampligase (Biosearch Technologies, A0102K), 1 U/μL RiboLock RNase inhibitor, 0.001 U/μL Phusion DNA polymerase (ThermoFisher, F530S), 0.4 U/μL RnaseH (QIAGEN, Y9220L)] at 37 °C for 20 min and then 45 °C for 45 min. We then performed the second RCA [1× Phi29 buffer, 0.2 mg/ml BSA, 0.25 mM dNTPs, 125 μM aminoally-dUTP, 1 U/μL Phi29 polymerase, 5% (vol/vol) Glycerol] overnight at 25 °C. We crosslinked the sample again at room temperature for 30 min and quenched crosslinking with 1 M Tris-HCl for 30 min.

We performed the BARseq3 Sequencing Module and imaged samples using a 25× Silicone oil objective (NA 1.05) on a Nikon TI2E microscope with a Crest Xlight V3 confocal unit, using 514 nm (G/T) and 640 nm (A/C) excitation lasers and acquired images on two Hamamatsu ORCA-FusionBT cameras synchronized via hardware TTL triggering. We used a ZT405/514/635rpc dichroic with a ZET532/640m emission filter, and a camera cube containing a T637/105dcbp-UT2 camera dichroic and a ET550/40+752/95m cleanup filter to separate the four sequencing channels. Briefly, we hybridized gene sequencing primer with FISH buffer [1 µM sequencing primer, 2× SSC (Invitrogen, AM9770), 10% formamide (Thermo Fisher, AM9344)]. We sequenced the gene and barcode signals using Illumina sequencing reagents from an MiSeq Reagent Nano 500 cycle Kit v2 (Illumina, MS-103-1003). For first round sequencing, we incubated samples in incorporation buffer at 60 °C for 3 min and washed the samples with PBSTw (2% Tween-20 in PBS) at room temperature. We then incubated samples in Iodoacetamide blocker solution [2.6 mg/mL Iodoacetamide (Thermo Fisher Scientific, A39271) in PBSTw] at 60 °C for 3 min. We washed samples again with PBSTw and followed by two washes with incorporation buffer. We then performed incubations in IRM at 60 °C twice for 3 min each. We then washed the samples four times with PBSTw at 60 °C for 3 min each. Then we imaged the gene signals. For all subsequent sequencing rounds, we washed samples twice in incorporation buffer and incubated samples in CRM at 60 °C for 3 min. We then followed the first-round sequencing procedure. We did Nissl staining (Invitrogen, N21479) with 1:20 dilution in PBST for 20 min after the third-round sequencing and imaged the Nissl signal at the third-round imaging cycle. After completing gene sequencing, we stripped the gene sequencing primer using stripping buffer (2× SSC, 80% formamide) at 60 °C three times for 3 min each and hybridized the barcode sequencing primer in FISH buffer at room temperature for 10-15 minutes. We imaged the Nissl signal again at the second barcode sequencing round.

### BARseq2 library preparation and *in situ* sequencing

We designed BARseq2 probes using the MATLAB script described in^19^. For each gene we designed 12 probes for a panel of 36 genes in marmoset and 167 genes in macaque. All the probed for genes and associated probe sequences are provided in Supplemental Table 2. We cryosectioned the samples at 20 µm thickness, mounted onto slides and stored slides in -80°C. We prepared BARseq2 libraries as previously described for marmoset and macaque virus screening^19^. Briefly, we removed slides from -80°C storage and immediately fixed in 4% paraformaldehyde (PFA) in PBS for 1h at room temperature. Following a PBS wash, we mounted HybriWell-FL chambers over the tissue sections. We dehydrated samples sequentially in 70%, 85%, and 100% ethanol for 5 min each, transferred to fresh 100% ethanol, and incubated for 1.5 hours at 4 °C. Sections were then rehydrated by repeated washes in PBST (PBS containing 0.5% Tween-20).

We used 50 μM random N20 primer (XC2757) and 1 μM barcode-specific LNA primer (XC1215) for endogenous mRNAs and Sindbis barcode RNA, respectively. We performed reverse transcription in 1× RevertAid RT buffer containing 20 U/μL RevertAid H Minus M-MuLV reverse transcriptase (Thermo Fisher Scientific), 500 μM dNTPs (Thermo Fisher Scientific), 0.2 μg/μL BSA (New England Biolabs), and 1 U/μL RiboLock RNase inhibitor (Thermo Fisher Scientific) at 37 °C overnight in a humidified chamber. The following day, we washed samples in PBST and crosslinked the cDNA using BS(PEG)9 [50 mM BS(PEG)9, 0.5% Tween-20 in PBS] at room temperature for 1 hour. We washed the samples once with 1M Tris-HCl, pH 8.0, followed by incubation in fresh 1M Tris-HCl for 30 min. We then washed samples twice with PBST (0.5% Tween-20 in PBS). We hybridized gene padlocks and followed by circularizing [1× Ampligase buffer containing 100nM of each padlock probe, 0.5 U/μL Ampligase, 0.4 U/μL RNase H, 1 U/μL RiboLock RNase inhibitor, an additional 50 mM KCl, and 20% formamide] at 37 °C for 30 min followed by 45 min at 45 °C. We then hybridized barcode padlocks and followed by gap-filling and circularizing [1× Ampligase buffer, 50 mM KCl, 0.1 μM barcode padlock, 0.05 mM dNTPs, 5% (vol/vol) Glycerol, 20% (vol/vol) formamide, 0.5 U/μL Ampligase, 1 U/μL RiboLock RNase inhibitor, 0.001 U/μL Phusion DNA polymerase, 0.4 U/μL RnaseH] at 37 °C for 5 min and then 45 °C for 45 min. We then washed twice with PBST and once with hybridization buffer (2× SSC, 10% formamide). We hybridized RCA primer (1 μM XC1417 in hybridization buffer) at room temperature for 15 min, followed by three 2-min washes in hybridization buffer and PBST washes.

We performed RCA [1× phi29 buffer, 0.25 mM dNTPs, 125 μM aminoallyl-dUTP, 0.2 μg/μL BSA, 5% glycerol, and 1 U/μL phi29 polymerase] at room temperature overnight. After RCA, we washed samples in PBST and crosslinked with BS(PEG)9 in PBST for 1 hour at room temperature. We quenched crosslinker with 1 M Tris-HCl, pH 8.0, for 30 min, followed by PBST washes.

We performed BARseq2 *in situ* sequencing using Illumina Miseq nano v2 kits, following previously established protocol^63^. We performed fluid exchange and image acquisition during sequencing either manually (Z13082) or using a home-made automated fluidics and imaging system, built on a Nikon TI2E microscope with a Lumencor Celesta laser, a Prion NanoScan OP400 piezo objective scanner, Crest Xlight v3, and Photometrics Kinetix cameras. We used the same filters and imaging conditions as described previously^63^.

### Carrier protein screening

We cryosectioned the samples into 300 µm using Leica CM 3050S cryostat at -12 °C. We melted the sections on the slide and quickly refroze them on the dry ice. We dissected brain regions (mouse: M1, thalamus, SNr, and medulla; zebra finch: HVC, RA, and Area X) on a metal block partially submerged in freezing 2.25 M CaCl2 (Sigma, 442909) solution^18^. We extracted RNA using Trizol reagent (Thermo Fisher, 15596026) following the manufacturer’s instructions. We then generated MAPseq libraries for all samples following the MAPseq2 protocol^32^. Briefly, we used reverse transcription primer with 12-bp random UMI sequences and 8-bp unique Sample specific identifier (SSI) for each sample (Primer sequence: 5’-GAC GTG TGC TCT TCC GAT CT -N12-X8-CAC GAC GGC AAT TAG GTA GC-3’, where X refers to SSI bases). We incubated samples with reverse transcription primer [4 µL RNA sample, 1 µL 10 mM dNTPs, 1 µL spike-in RNA (10^5^ molecules), 1 µL 10 µM RT primer, 9 µL ddH2O] at 65 °C for 10 min and immediately placed samples on ice for 1 min. We then incubated samples with reverse transcription reaction mix [4 µL 5× MashUp buffer,1 µL DTT (100 mM), 1 µL SUPERase In RNase Inhibitor (Thermo Fisher, AM2696), 1 µL MashUp RT enzyme] at 50 °C for 60 min. We treated each sample with 2 µL Thermolabile Exonuclease I (New England Biolabs, M0568S) at 37 °C for 15 min and followed by inactivation at 80 °C for 10 min. We then treated sample with 2 µL RNase If I (New England Biolabs, M0243S) at 37 °C for 60 min. We cleaned the sample with AMPure XP bead. We then performed PCR1 with 5× reaction mix [5× SYBR green I (Invitrogen, S7563), 5× AccuPrime reaction mix (Invitrogen, 123444024), 2.5 mM MgSO4, 5 μM foward primer (Primer sequence: 5’-CGAGAAGCGCGATCACATG -3’), 5 μM reverse primer (Primer sequence: 5’-CTGGAGTTCAGACGTGTGCTCTTCCGATCT-3’), 10% ROX Reference Dye (Invitrogen,12223012), and 25% DMSO] and stopped the reaction during exponential phase of PCR to avoid saturation. We diluted the PCR1 product about 100-fold and performed PCR2 for 6-10 cycles with unique i5 and i7 Illumina indexing primers. After PCR2, we gel extracted the 219 bp PCR with the QIAquick kit (Qiagen, 28704), and measured its concentration using a TapeStation (Agilent, G2992AA). We then pooled the libraries and sequenced them on the Illumina NovaSeq X plus platform using paired-end sequencing.

### Zebra finch HVC projection mapping

We dissected HVC, RA, Area X and cerebellum tissue as described for carrier protein screening and then prepared the MAPseq2 libraries as described above.

### *In situ* sequencing data processing

#### BARseq3

We used a custom computational pipeline to process BARseq3 image data as previously described^20^. Briefly, we performed maximum z-projection and aligned imaging channels using a homography transform and registered each round to the round containing the Nissl signal by using an affine transform for coarse alignment followed by Demons alignment for nonlinear refinement. We then performed color bleed correction, min–max normalization, and background subtraction (radius = 5). Due to the small number of genes probed for, the min-max normalization parameters for each channel were determined in a sequencing round containing a lot of signals and then applied to those channels across all rounds to avoid the background signal from being artificially amplified in rounds where that channel contained relatively few signals. For gene signal, we detected amplicons using Bardensr^64^ and assigned confidence scores to candidate gene IDs based on matches to the gene codebook. We used a confidence threshold of 0.75 for gene call. We used Cellpose 4^65^ to segment cells based on Nissl signal and assigned gene calls to cells based on spatial overlap with the corresponding cell masks to generate the expression matrix. To process barcode data, we aligned barcode images first and segment cells based on Nissl signal in barcode rounds using Cellpose 4^65^. We aligned the Nissl signal masks in barcode and gene rounds and applied the transformation to barcode images. We defined soma barcode signal as a region in which the intensity of the strongest channel was at least three times that of the weakest channel and the signal-positive area covered more than 40% of the Nissl mask. We assigned the nucleotide at each sequencing cycle based on the maximum intensity nucleic-acid fluorescence signal within each cell mask and excluded cells with a barcode mask area < 700 pixels. Finally, we mapped the spatial coordinates of barcoded cells onto the gene signal cell spatial location to link each barcoded cell to its corresponding gene-expression profile.

### BARseq2

We processed BARseq2 *in situ* sequencing images using a MATLAB pipeline as previously described^19^. Briefly, we denoised maximum-intensity projections with Noise2Void^66^, corrected for channel crosstalk and background, and registered across sequencing and hybridization cycles. We stitched fields of view into whole-section images using MIST^67^, and identified gene rolonies and base-called with Bardensr^64^ against the padlock codebook. We segmented cells with Cellpose^68^ and dilated masks by 3 pixels to accommodate residual registration error, and assigned rolonies within each mask to the corresponding cell to generate the cell-by-gene count matrix. We removed duplicate cells in overlapping fields of view by spatial matching within 10 µm and registered barcode images to the gene-expression images through a shared reference channel. We called soma barcodes from the registered barcode images using two approaches. First, we averaged channel intensities across all pixels within each cell mask and assigned each cycle to the brightest channel. Second, we made calls using only high-purity pixels: pixels were retained when their third lowest per-cycle purity exceeded 0.85, and cells containing more than 50 such pixels were called from their mean channel intensities. We used the high-purity-pixel call when available; otherwise, we used the whole-mask call. We classified soma as barcoded when the called sequence passed a sequence-complexity threshold and both the third lowest per-cycle purity and third-lowest maximum-channel intensity exceeded 0.85 and 150, respectively. This automated calling was followed by manual annotation to remove cells that were mis-called as barcoded because of strong barcode labeling in neighboring cells and/or passing axons and/or dendrites. Using the third lowest value allowed up to two sequencing cycles with poor signal without rejecting an otherwise high-quality barcode.

### Defining cell types

#### BARseq3

In mice, genes with fewer than three counts per cell were excluded to minimize potential signals originating from axons or dendrites. A cell was classified as an excitatory neuron if Slc17a7 had the highest count and the Gad1/Slc17a7 count ratio was less than 0.67; otherwise, it was classified as unknown. Similarly, a cell was classified as an inhibitory neuron if Gad1 had the highest count and the Slc17a7/Gad1 count ratio was less than 0.67; otherwise, it was classified as unknown. Cells in which GFAP had the highest count were classified as astrocytes. Cell types in zebra finches were classified using the same strategy. Slc17a6 and Gad1 were used to identify excitatory and inhibitory neurons, respectively, whereas PLP1, SOX9, and Csf1r were used to identify glial cells. In frogs, Snap25 signals with fewer than three per cell were excluded, whereas Slc1a2 and Slc4a4 signals with fewer than two counts per cell were excluded. A cell was classified as a neuron if Snap25 had the highest count and the ratio of the higher count between Slc1a2 and Slc4a4 to the Snap25 count was less than 0.67; otherwise, it was classified as unknown. Conversely, a cell was classified as an astrocyte if either Slc1a2 or Slc4a4 had the highest count and the ratio of the Snap25 count to the higher count between Slc1a2 and Slc4a4 was less than 0.67; otherwise, it was classified as unknown.

### BARseq2

We resolved cells type identities as previously described^63^. We excluded cells with fewer than 20 total gene counts or fewer than 5 detected genes, and normalized gene counts to 10 counts per cell and log2-transformed normalized values. We performed Clustering iteratively as previously described^63^. At each round, we computed the first 30 principal components, constructed a shared-nearest-neighbor graph (k = 15; Jaccard edge weighting; approximate nearest neighbors), and partitioned the graph by Louvain community detection; UMAP (k = 15) was used for visualization. In the first round, we separated glutamatergic, GABAergic, and non-neuronal cells. In the second round, we analyzed the glutamatergic and GABAergic populations separately to identify subclasses. In the third round, we analyzed each glutamatergic subclass independently to identify clusters. At each round, we annotated clusters based on enrichment of curated marker genes and excluded clusters showing poor transcript detection rather than a distinct biological identity from cell-type analyses.

### MAPseq data analysis

#### Producing the raw barcode matrix

To generate the barcode matrix from raw sequencing reads, we followed MAPseq2 pipeline^32^. Briefly, we extracted 28 nt of the barcode sequence (26 nt barcode sequence for carrier protein screening) from Read 1 and 20 nt sequence (12 nt UMI and 8 nt SSI) from Read 2 FASTQ files. We filtered reads based on the correct SSI sequences and removed sequences containing undetermined nucleotides. We then trimmed off the SSI sequences, retaining only the barcode and UMI sequences. We generated a rank plot for each sample based on abundance of each read using MATLAB (MathWorks, R2022a) and discarded low abundance, low confidence reads in the tail of the rank plot^69^. We separated barcodes from spike-ins based on spike-in specific sequence end to the barcode (CGTC). We then used bowtie^70^ to do an all-to-all alignment, and collapsed barcode sequences with up to 3 mismatches to the most abundant sequence to correct for sequencing and PCR errors. Finally, we constructed the final barcode matrix by matching barcodes across samples using MATLAB.

### Carrier protein screening

For carrier protein screening, we identified barcodes carried by each carrier protein based on the carrier protein specific identifier sequence. We filtered the barcode matrix by requiring at least 50 UMI counts in the injection site, at least 3 UMI counts in the strongest target region and removed the counts less than 2, and also removed all barcodes for which the UMI count in the injection site was greater than maximum UMI count in target regions. For each target region, we removed the carrier protein if the total number of barcode sequences in that region was lower than 3. Cumulative distribution functions were generated with a bin size of 50 UMIs. We ranked the carrier proteins based on their mean barcode counts within each brain region or sample. We then used the average value of the ranks across samples to plot the relative performance of each carrier protein.

### HCV projection mapping

We filtered the barcode matrix by requiring at least 75 UMI counts in the injection site, at least 5 UMI counts in the strongest target region, and removed all counts less than 2, and also removed all barcodes for which the UMI count in the injection site was 5 times greater than maximum UMI count in target regions. We first normalized barcode UMIs to the spike-in RNA and then normalized the projection matrix row-wise by dividing each element by the sum of all elements in each row. For comparison, we obtained the per neuron binarized projection data of published single-neuron HVC reconstructions, and built a projection matrix by scoring projections to RA and Area X as present or absent.

## MATERIALS AVAILABILITY

All plasmids created for this study are available on Addgene under accession numbers #262062 to #262102. A reference table is provided as Supplemental Table 3.

## DATA AND CODE AVAILABILITY

Analysis code and processed data files are available at GitHub (https://github.com/huihqi/Universal_MAPseq) and Zenodo (DOI:10.5281/zenodo.21997210). All raw sequencing datasets are available in the NCBI Sequence Read Archive (SRA) under accession number PRJNA1513960. Processed MAPseq data for zebra finch HVC are also provided as Data S1.

## ACKNOWLEDGMENTS

We thank Diane E. Griffin and Michael S. Diamond for generously providing us with plasmid constructs and Kebschull lab members for valuable feedback on this manuscript. This work was supported by NIH BRAIN Initiative grant R34132027 [J.M.K., M.A.L.], BRAIN CONNECTS grant U01NS13216 [J.M.K., X.C., G.H., P.H.R.], NIH grant ZIAMH002984 [J.B.], and NIH grant R01NS075044-11 [M.A.L.]; the Packard Fellowship, Sloan Fellowship, and Klingenstein Simons Fellowship to J.M.K.; a MITACS travel fellowship to X.W.; a Kavli Distinguished PhD fellowship to M.M-G.A.; an NSF GRFP and NIH 5T32NS091018-24 support to C.S.. The content is solely the responsibility of the authors and does not necessarily represent the official views of the National Institutes of Health.

## AUTHOR CONTRIBUTIONS

H.Q. and J.M.K. conceptualized the study; H.Q. performed most of the experiments and data analyses; X.W. tested non-mutated viruses in zebra finches, performed HVC injections, and collected the tissue with support from J.T.S.; C.S. generated MAPseq libraries for HVC tracing experiments; Z.Y. performed parrot and cowbird injections with support from M.A.L; J.C. tested pseudotyped viruses in the mouse cerebella nuclei; M.M. and C.E. performed virus screening in macaques with support from G.D.H., P.H.R. and X.C.; A.F. performed virus screening in marmoset with support from J.B.; C.W. generated carrier protein screening library; M.M-G.A. and D.Z.F-G. provided assistance with BARseq3 data analysis and troubleshooting.; M.L. assisted with animal husbandry and breeding; H.Q. and J.M.K. wrote the manuscript with feedback from all authors; J.M.K. supervised the project.

## Supplementary Figures

**Supplementary Figure 1.**
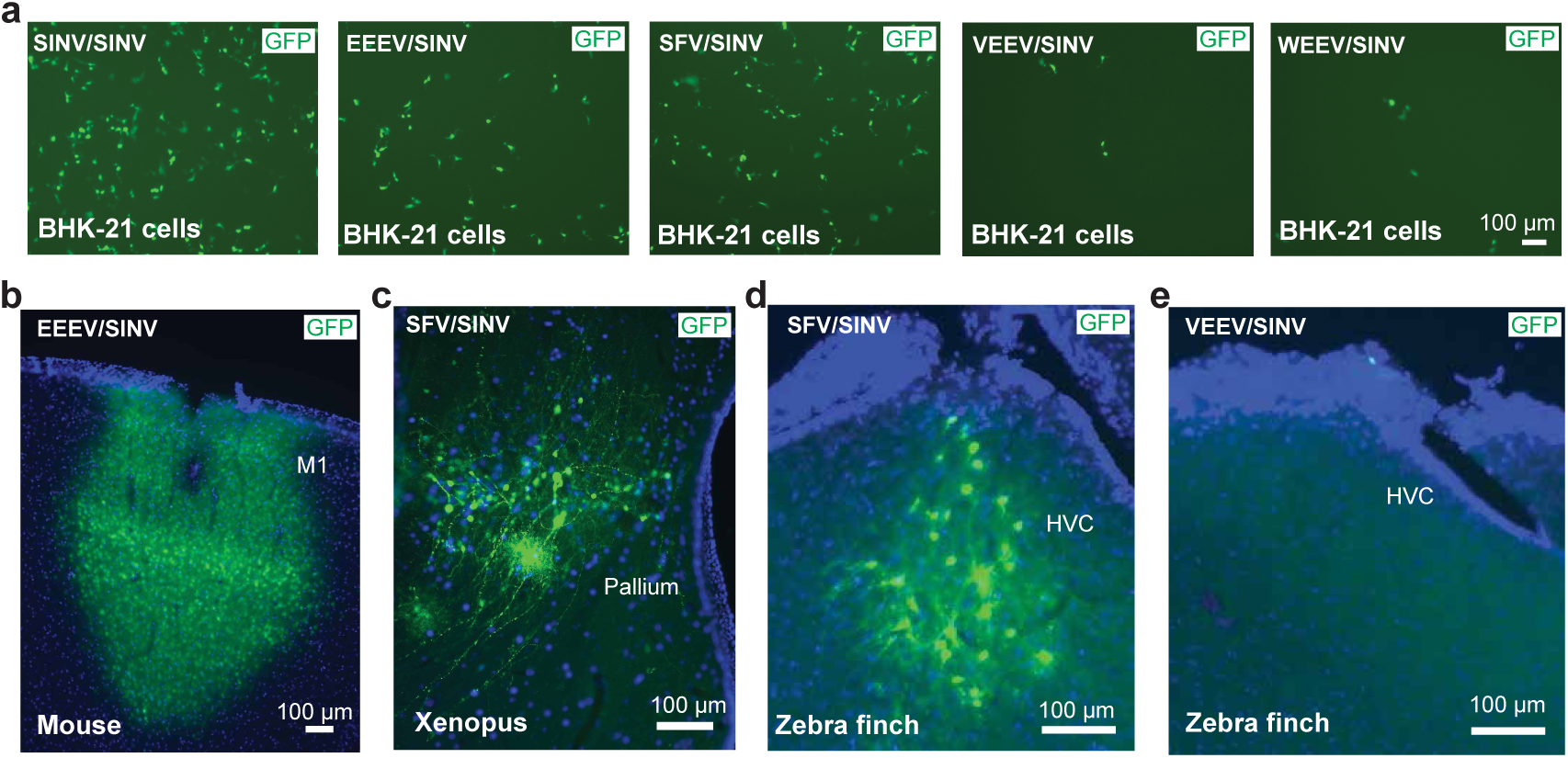
Pseudotyped viruses exhibit species-and cell type-dependent differences in infectivity. (a) SINV/SINV, EEEV/SINV, and SFV/SINV show robust infection in BHK-21 cells, whereas VEEV/SINV and WEEV/SINV show limited infection. Cells were infected with the same number of viral particles, as measured by qPCR of RNAseI protected genomes. (b-e) EEEV/SINV and SFV/SINV show robust infection across mouse M1, Xenopus pallium, and zebra finch HVC, whereas VEEV/SINV fails to infect zebra finch HVC, consistent with the screening results.

**Supplementary Figure 2.**
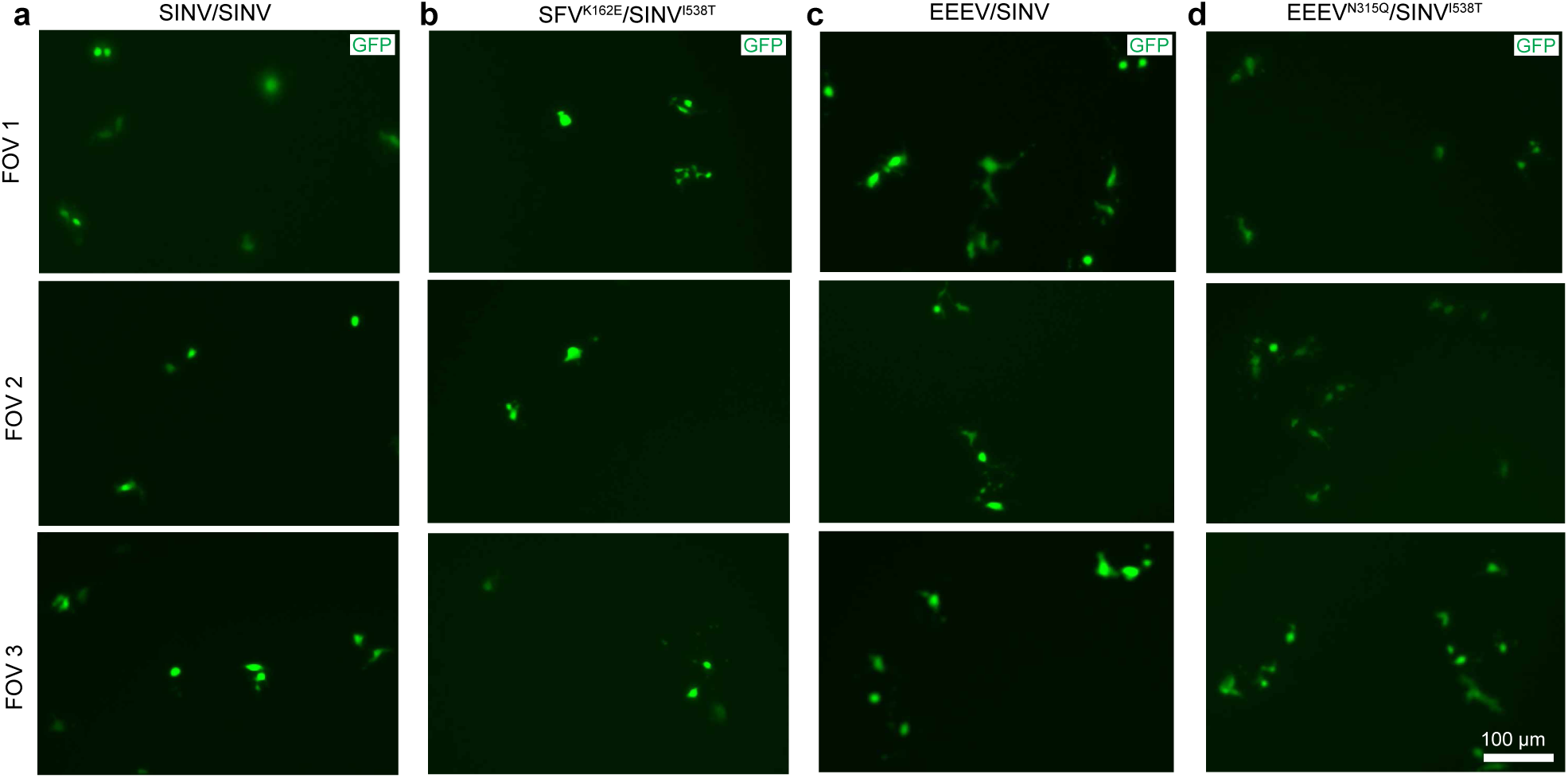
Pseudotyped viruses are propagation incompetent. (a-d) Plaque assay in BHK-21 cells showing that the original MAPseq (SINV/SINV), SFV^K162E^/SINV^I538T^, EEEV/SINV and EEEV^N315Q^/SINV^I538T^ virus do not produce plaques as expected for propagation incompetent viruses. Only individual infected cells are visible.

**Supplementary Figure 3.**
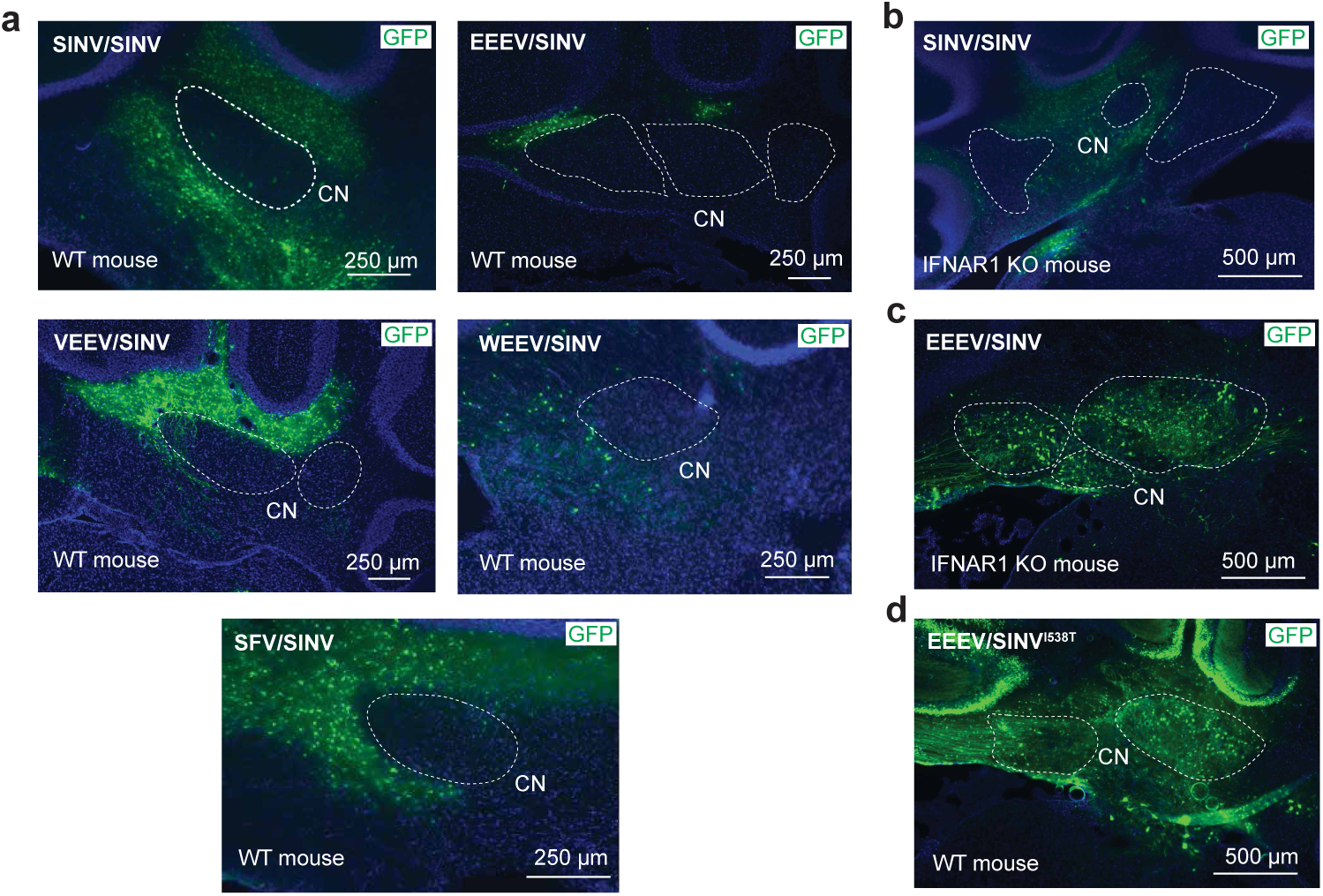
Attenuation of host innate immune defenses enables robust infection of mouse cerebellar nuclei. **(a)** Conventional pseudotyped viruses show no or little infection in mouse cerebellar nuclei (CN). (b, c) EEEV/SINV shows robust infection in the cerebellar nuclei of IFNAR1 knockout mice (c), whereas SINV/SINV remains ineffective (b), indicating that both viral structural proteins and interferon signaling play a key role in enabling infection. (d) EEEV/SINV^I538T^ overcomes innate immune restriction and produces robust infection in the cerebellar nuclei of wild type mice.

**Supplementary Figure 4.**
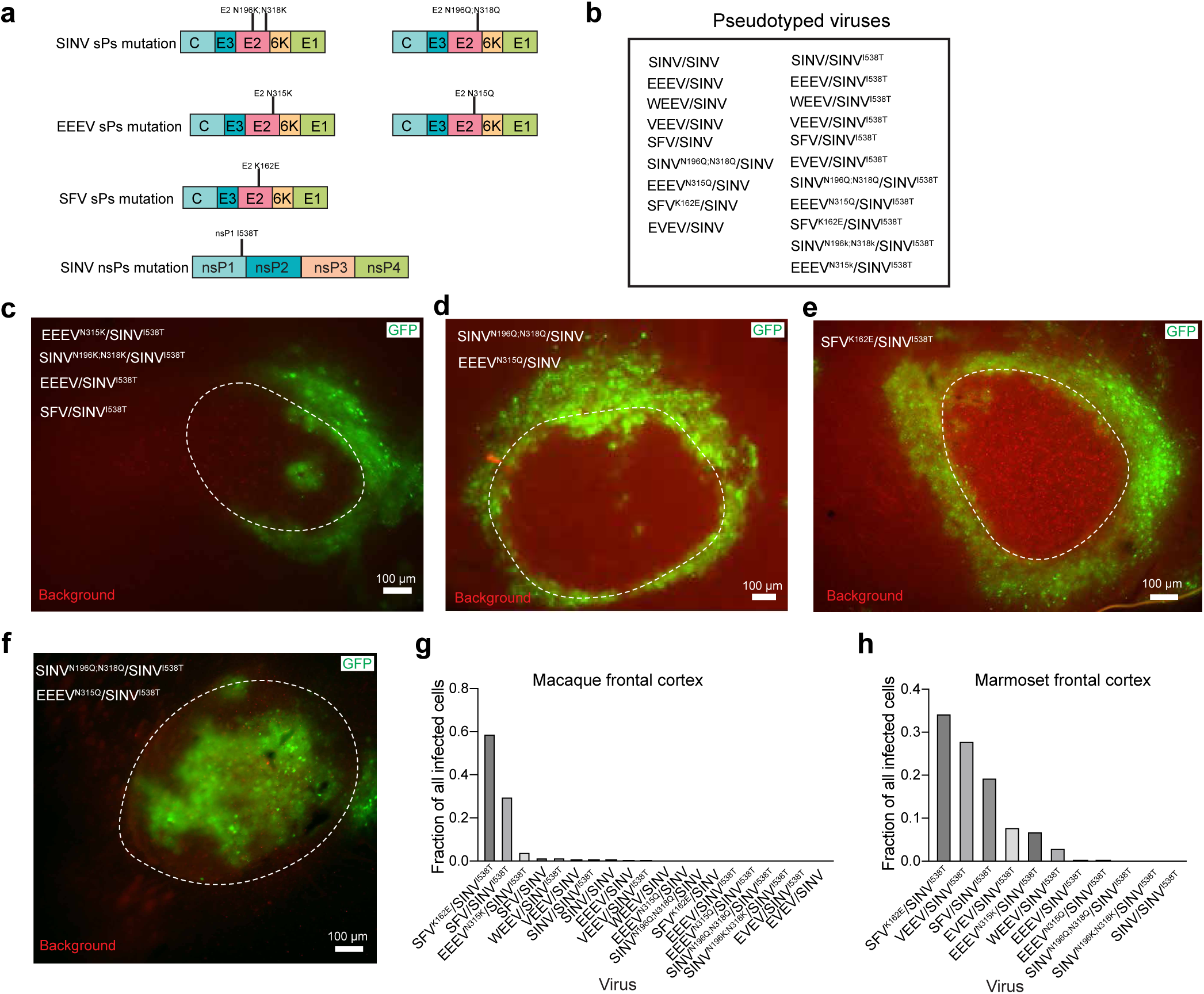
Screening engineered viruses in zebra finch, macaque, and marmoset. (a) Locations of point mutations in viral structural proteins and the non-structural proteins. (b) Combinations of all structural proteins and viral genomes used for screening. (c-e) EEEV^N315K^/SINV^I538T^, SINV^N196K;N318K^/SINV^I538T^, EEEV/SINV^I538T^, SFV/SINV^I538T^, SINV^N196Q;N318Q^/SINV, EEEV^N315Q^/SINV, and SFV^K162E^/SINV^I538T^ show little or no detectable infection in zebra finch RA. (f) A mixture of SINV^N196Q;N318Q^/SINV^I538T^ and EEEV^N315Q^/SINV^I538T^ produces robust infection in zebra finch RA. (g, h) Screening of engineered viruses in macaque and marmoset frontal cortex, showing the infection results for all viruses in the pool. SFV^N315Q^/SINV^I538T^ shows robust infection in both species.

**Supplementary Figure 5.**
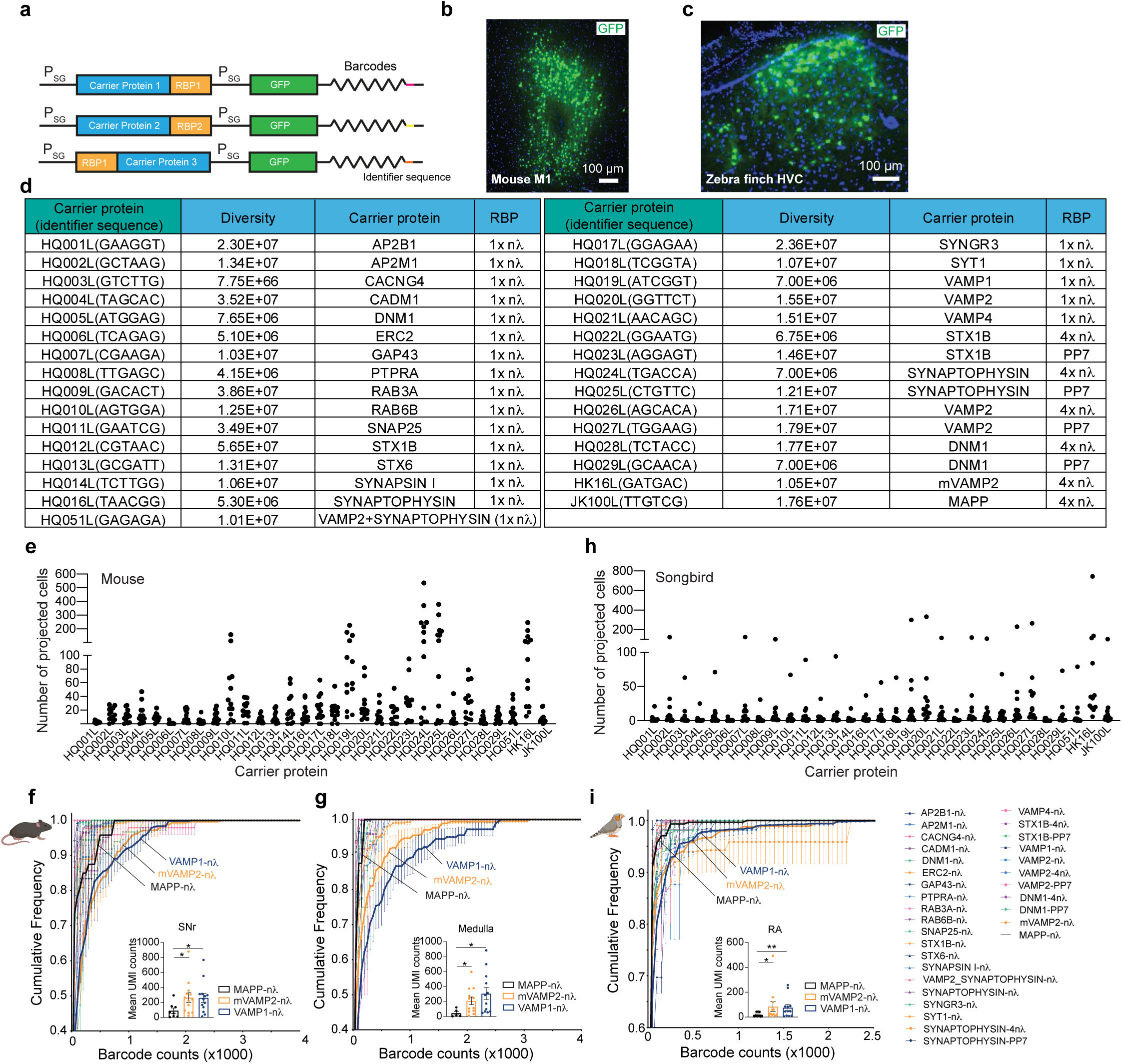
Screening carrier proteins in mouse and zebra finch. (a) Genome design used for carrier protein screening. [RBP: RNA-binding protein (1× nλ, 4× nλ, or PP7)]. (b, c) Infection of pooled carrier protein libraries in mouse M1 (b) and zebra finch HVC (c). (d) Information on the carrier protein plasmid libraries, including barcode diversity. (e) Number of projecting neurons recovered for each carrier protein in mouse M1 (n = 13 hemispheres). (f, g) Carrier protein screening results from mouse M1, shown as the cumulative distribution of barcode UMI counts for each carrier protein in the substantia nigra pars reticulata (SNr) and medulla. VAMP1-nλ and mVAMP2-nλ are the top-performing candidates, both outperforming MAPP-nλ. Insets show mean UMI counts in each target region for the MAPP-nλ control and the two top performers. (h) Number of projecting neurons recovered for each carrier protein in zebra finch HVC (n = 12 hemispheres). (i) Carrier protein screening results from zebra finch HVC, shown as the cumulative distribution of barcode UMI counts for each carrier protein in RA. VAMP1-nλ and mVAMP2-nλ are the top-performing candidates, both outperforming MAPP-nλ. Insets show mean UMI counts in each target region for the MAPP-nλ control and the two top performers (Mouse: n = 13 hemispheres, Zebra finch: n = 12 hemispheres, Mann-Whitney test, * p<0.05, **p<0.01).

**Supplementary Figure 6.**
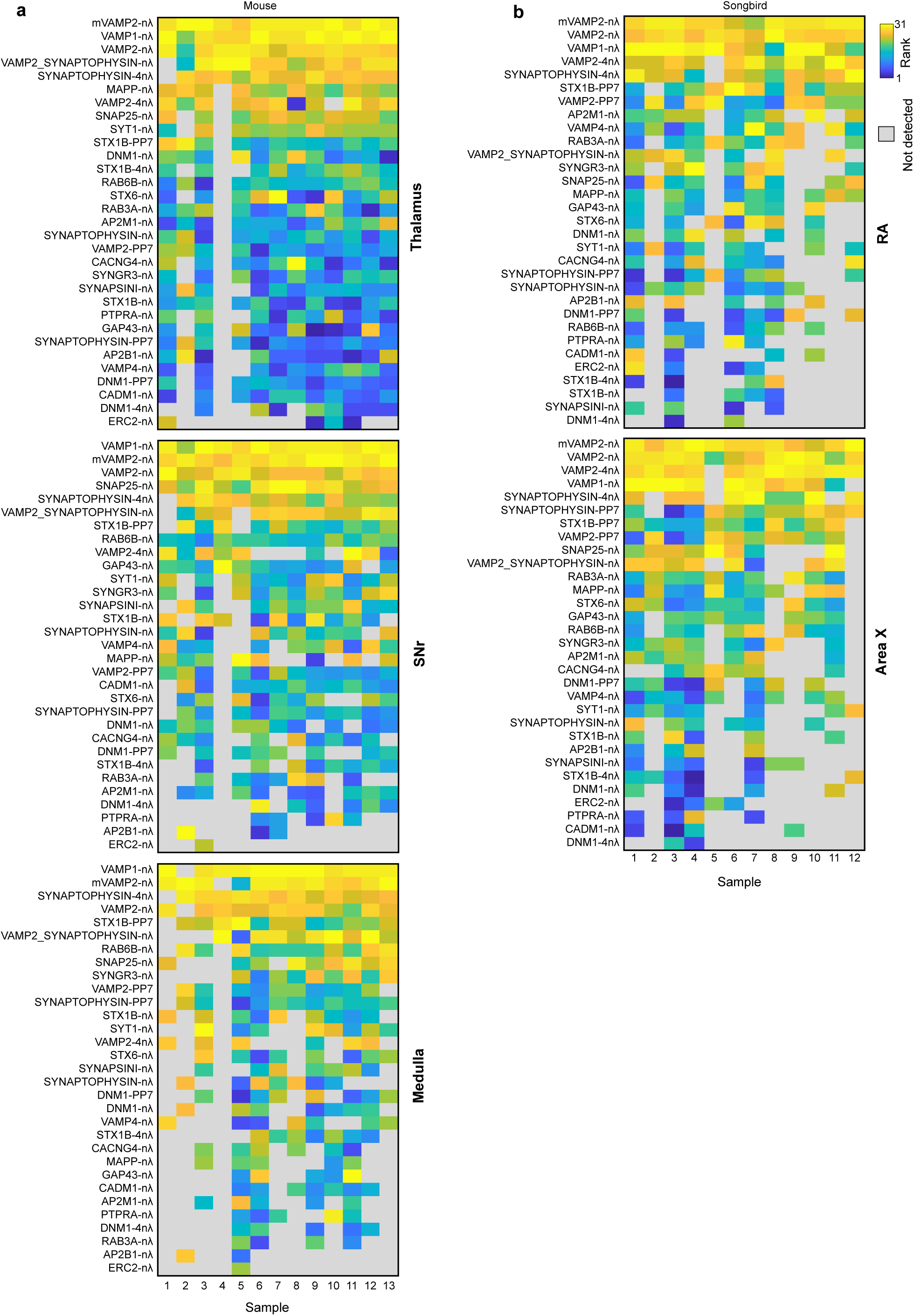
Ranking of carrier proteins in individual samples and brain regions. **(a, b)** Heat maps of carrier protein performance ranks measured by mean barcode UMI counts show the consistently high rankings of VAMP1-nλ and mVAMP2-nλ across samples and brain regions.

**Supplementary Figure 7.**
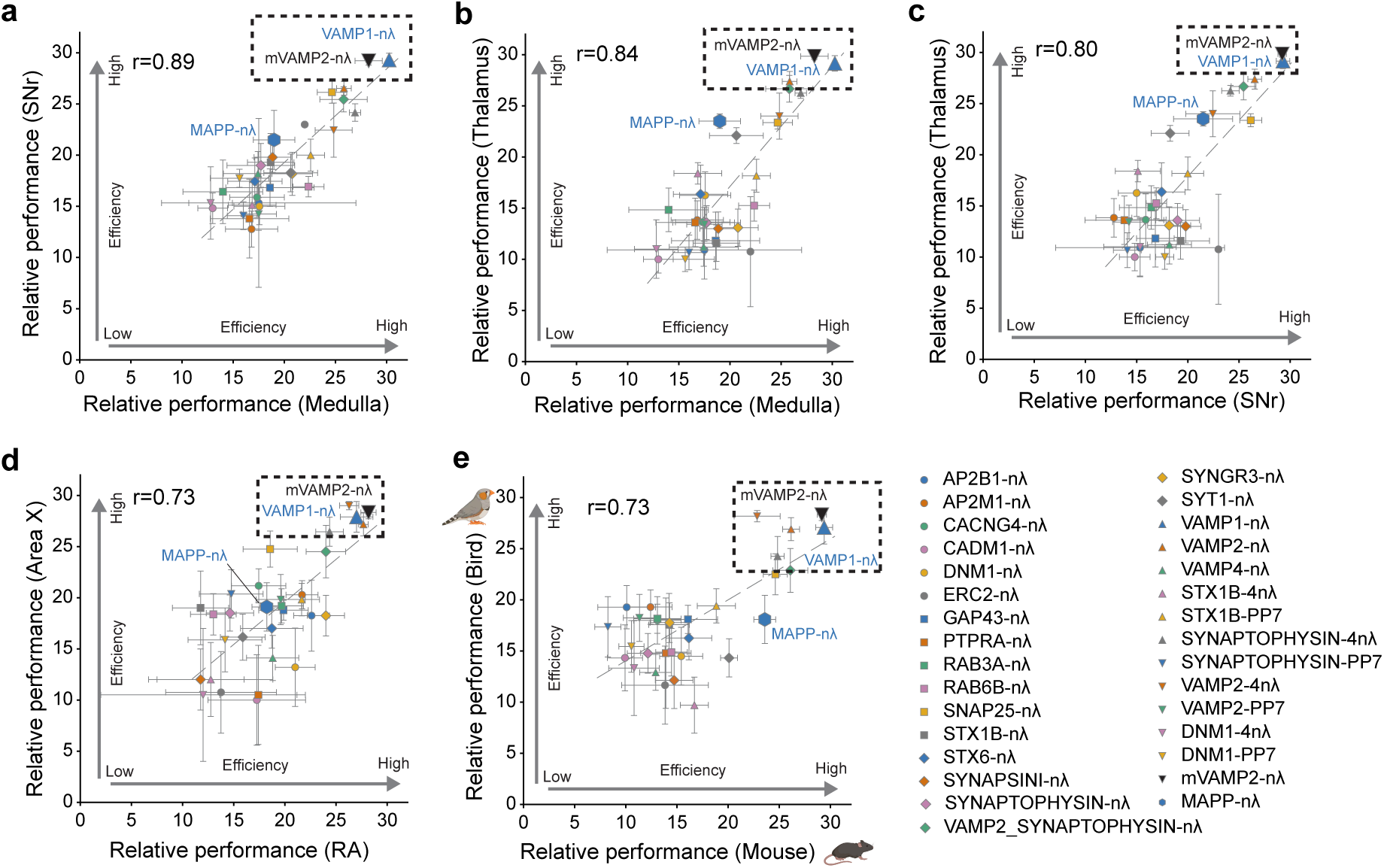
Correlations of carrier protein ranks across brain regions and species. **(a-e)** Carrier protein rankings were highly correlated across brain regions (Thalamus, SNr, Medulla, Area X and RA) and species (mouse and zebra finch).

**Supplementary Figure 8.**
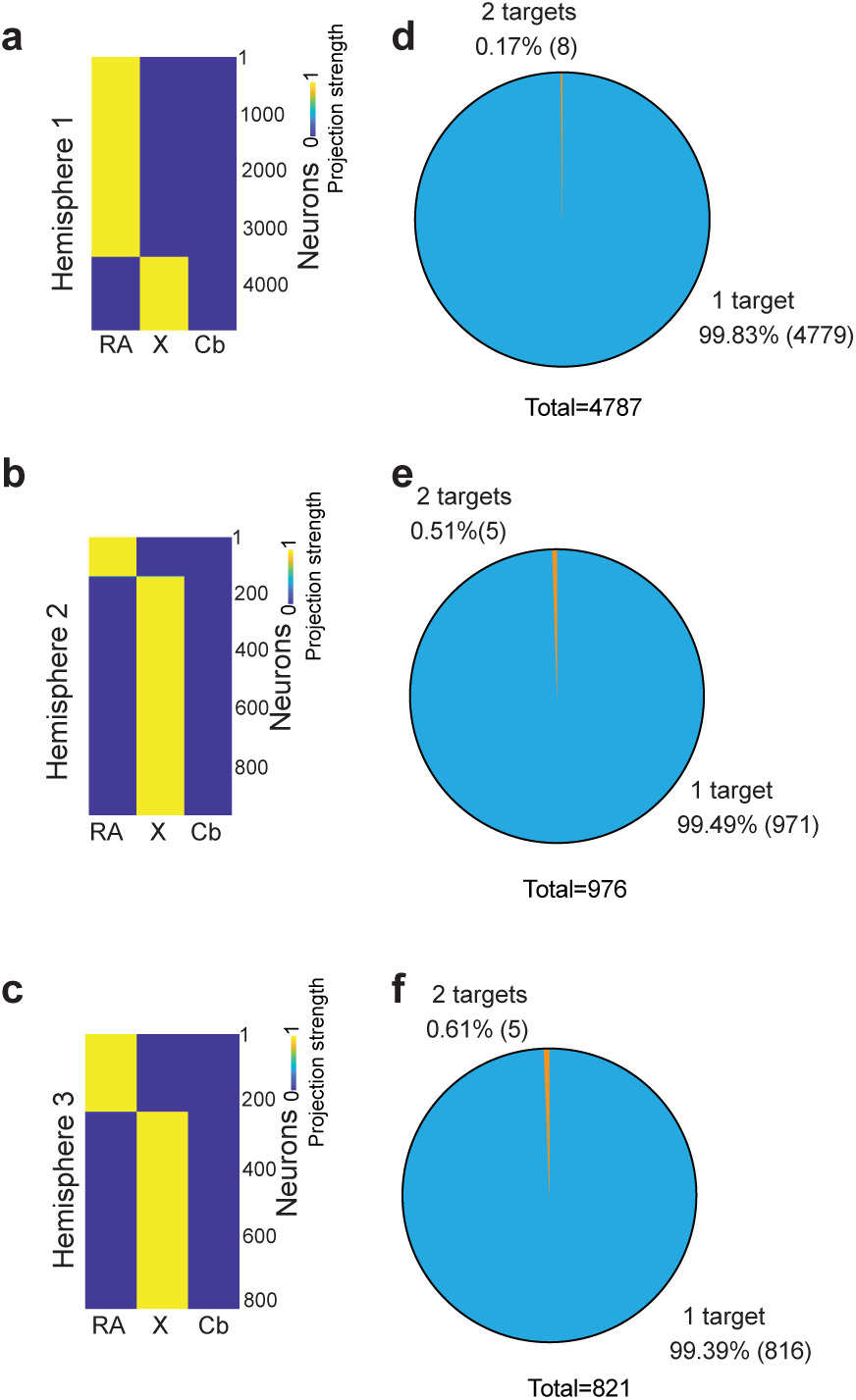
Projection patterns mapped in individual zebra finch HVC injections. **(a-c)** Projection matrices of HVC neurons from each mapped hemisphere, showing robust projection mapping in each biological replicate. Hemispheres 2 and 3 are from the same animal. (d-f) Pie charts showing the number of target regions per neuron in each hemisphere.

