## Supplementary Data S1 for "Universal barcoded connectomics for massively parallel single-neuron projection mapping in diverse vertebrate species": README.rtf

file 49L: Songbird49 left hemisphere
file 49R: Songbird49 right hemisphere
file 50R: Songbird50 right hemisphere


Data in the .mat files:

B: injection and target sites barcode counts
Bnorm: normalized barcode counts
B_tar: target sites barcode counts
Bnorm_tar: normalized target sites barcode counts

Column order:
B and Bnorm: HVC, RA, X, Av, Cb
B_tar and Bnorm_tar: RA, X, Av, Cb
